# Evolution of informed dispersal strategies in trophic meta-communities

**DOI:** 10.64898/2026.07.17.739253

**Authors:** Yuriy Pichugin, Corina Tarnita

## Abstract

Living organisms move and their movement is both a response to local conditions and, often, the cause of change in those conditions. This feedback loop is often overlooked in theoretical studies on the evolution of dispersal, which assume that either the decision to leave is uninformed or that the local conditions are exogenously driven. Here, we embrace the feedback loop and study what dispersal strategies evolve in a trophic meta-population where the dynamics are entirely endogenous. We show that there are five possible classes of strategies that can evolve depending on the ecological conditions: leaving once local conditions fall low enough (*Unsaturated*), having an extended stay even under adverse conditions (*Saturated*), leaving from a high quality location (*Anxious*), leaving independently of the location state (*Ignorant*), and completely abstaining from dispersal (*No-dispersal*). The *Unsaturated* class captures the classical prediction of the marginal value theorem, while the other four extend the range of possible evolutionarily optimal strategies. Which class of strategies evolves depends on the kind of information being sensed (resource availability versus conspecifics density), the size of the local consumer population at equilibrium, and the stability of this equilibrium. Our results provide a theoretical underpinning for the diversity of movement strategies observed in nature that deviate from classic predictions and suggest a comparative framework that can inform experimental design.

## 1 Introduction

Biological dispersal is defined as movement that creates a gene flow [Ronce, 2007], typically driven by the search for food, escaping predation, or the search for mates. Among these, foraging dispersal—dispersal driven by the search for food—has been widely studied for nearly a century, both empirically and theoretically. Yet, fundamental questions remain open, in great part because the evolution of dispersal strategies is embedded into an eco-evolutionary feedback loop, the complete study of which has remained analytically and computationally intractable: an adaptation of individuals’ strategy changes the population dynamics, which in turn, alters environmental conditions experienced by the species, thereby changing the selection pressure applied to the population, which may then challenge the original adaptation itself.

Valuable insights have been obtained regarding foraging dispersal by making several simplifying assumptions, such as studying the question as an optimization problem for one focal individual [Charnov, 1976], assuming that individuals have complete information about the entire environment [Fretwell and Lucas, 1969], or making the environmental state independent of the distribution and actions of dispersing populations [Parvinen, 2002, Gyllenberg et al., 2002, Parvinen et al., 2003, Poethke et al., 2003, Massol et al., 2011, Parvinen et al., 2020, Ohtsuki et al., 2020]. The most commonly considered dispersal strategy arising from these simplified models can be summarized as: ‘An individual should leave its current location only if the expected gains from an arbitrary alternative location are higher than from the current one; otherwise it should stay where it is.’ This strategy has been shown to be optimal in an environment with an infinite number of patches, where the focal individual has knowledge of the marginal gains acquired from moving away from its current patch (described as the marginal value theorem or MVT [Charnov, 1976]). The same strategy also regularly arises in single species meta-population models with local information and externally driven environmental states, where it is known as the “bang-bang” strategy [Gyllenberg and Metz, 2001, Metz and Gyllenberg, 2001, Kisdi, 2004]. At the scale of the population, the dynamics resulting from individuals employing such a strategy is expected to converge to the ideal free distribution (IFD) [Fretwell and Lucas, 1969], in which individuals distribute themselves so that everyone secures the same share of available resources and, once this state is reached, dispersal halts.

Although such predictions of simplified models found some empirical support [Pyke, 1978, Milinski, 1979, Harper, 1982, Milinski, 1984, Godin and Keenleyside, 1984, Cuthil et al., 1994, Wajnberg et al., 2000], there is also a growing body of examples that violate predicted patterns [Barash, 1974, Tamisier, 1974, Gaines and McClenaghan, 1980, Goss-Custard et al., 1984, Lessells, 1985, Parker and Sutherland, 1986, Stiling, 1987, Walde and Murdoch, 1988, Cassini et al., 1990, Crowley et al., 1990, Nonacs, 2001, DiGiorgio et al., 2020, O’Bryan et al., 2020, Sappington, 2024]. To address these discrepancies, various model extensions have been suggested, such as introducing predation risk or the search for mates in addition to foraging [Nonacs, 2001], considering individual heterogeneity in the ability to search for food, or compete with others [Whitham, 1980, Parker and Sutherland, 1986, Houston and McNamara, 1988], assuming externally driven local extinction [Parvinen, 2002, Gyllenberg et al., 2002], or erroneous evaluation of the environment [Kun and Scheuring, 2006, Poethke et al., 2016]. It is problematic, however, to know when it is necessary to incorporate additional factors to explain discrepancies if a true baseline of the main process has not been established. In other words, are the discrepancies the result of omitting additional processes (such as predation risk and mate search) or finer details (e.g., individual variation) or are they the result of working with a limited set of predictions resulting from oversimplified descriptions of the main process of foraging dispersal? To establish a baseline, dispersal has to be explored in its broadest eco-evolutionary context, where each adaptation is both a consequence of the applied selection pressures and the cause of further ecological change [Starrfelt and Kokko, 2012]. The study of a self-contained eco-evolutionary feedback loop has been attempted—especially in meta-communities models that feature the interaction of multiple species in a meta-population—but computational and analytical limitations have forced other types of simplifications [Gross et al., 2020]. Specifically, such models take a limited view of dispersal decisions by assuming that dispersal is completely uninformed (i.e. it uses neither the state of the current patch, nor information about the rest of the environment) [Pillai et al., 2012, Amarasekare, 2016, Chaianunporn and Hovestadt, 2019]. The main finding of such models is that the more likely the patch extinction, the higher the evolved dispersal rate [Pillai et al., 2012, Travis et al., 2013, Laroche et al., 2016]. A notable departure from these simplifying assumptions is [Khattar et al., 2024], who allowed for the possibility of informed dispersal in a multi-species, lattice-structured meta-community setting and found new classes of emerging dispersal strategies that showcase the nuance that can be obtained from considering non-constant dispersal functions. For instance, they, counterintuively, find the evolution of high dispersal rates from high quality patches. These results arise despite the computationally simplifying assumption of a one-parametric decision function (i.e. a power law) and a sparse sampling of ecological conditions, thus reinforcing the expectation that a richer baseline can be obtained from studying the evolution of informed dispersal in meta-communities [Gross et al., 2020, Fronhofer et al., 2024, McPeek et al., 2024].

Here, we present a model for the evolution of informed dispersal in a consumer-resource metacommunity, where we allow a complete eco-evolutionary feedback loop and systematically explore the space of possible ecologies. Furthermore, we consider a rich family of decision functions that includes previously considered functional forms (e.g., constant or concave). We circumvent computational challenges by employing a constructive classification approach informed by analytical results from the single-patch consumer-resource model. This allows us to explore both a much richer ecological space and a much richer decision space than all prior studies and, therefore, to contextualize prior findings within a broader baseline.

## 2 Model

### Population

We study a meta-population of *M* identical patches, inhabited by two species: a non-dispersing resource (*r*) and a potentially dispersing consumer (*c*), whose evolved dispersal strategies constitute our object of study. Consumers could be either in a foraging state (*f*), during which they perform resource acquisition or dispersal, or in a processing state (*p*), during which consumers eat the acquired resource and reproduce, see Fig. 1A. Dispersing consumers randomly select a patch to disperse to (i.e. the *M* patches form a complete graph).

**Figure 1:**
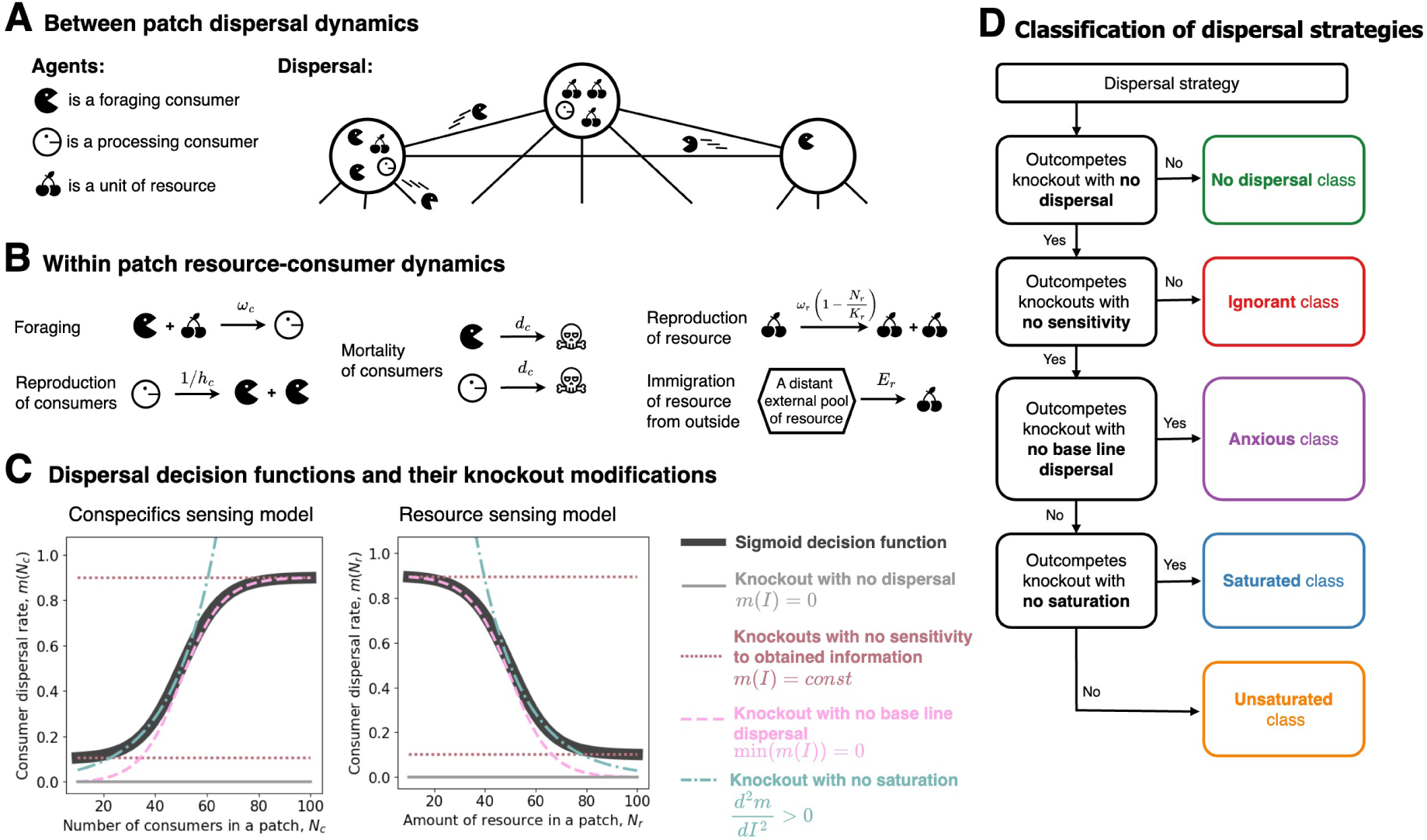
Model schematics. **A** Consumer and resource species inhabit a fully connected graph of patches forming a trophic meta-community, where foraging consumers are able to disperse between patches. **B** The population dynamics within a patch is determined by five processes: foraging (with rate *ω_c_*), reproduction (1*/h_c_*), and mortality (*d_c_*) of consumers, plus reproduction (*ω_r_* (1 *− N_r_/K_r_*)) and immigration of the resource from outside (*E_r_*). **C** The dispersal strategy is determined by a sigmoid decision function (black lines) of the sensed information (number of consumers *N_c_* or amount of resource *N_r_*), which parameters are optimized in the course of evolution. In both informed dispersal models, the performance of the evolved dispersal strategy (black) is tested in competitions against five “knockout” strategies: one does not disperse at all (solid gray line), two have no sensitivity to acquired information (dotted brown), one has no base line dispersal (dashed pink), and one has no saturation at detrimental conditions (dash-dotted turquoise). **D**: Classes are defined in their competitive ability against modified “knockout” strategies.

### Within-patch ecology dynamics

In the absence of consumers, the resource grows logistically (with growth rate *ω_r_* and carrying capacity *K_r_*). Consumers follow a predator-prey dynamics with Holling type II functional response. Foraging consumers must capture resources (at rate *ω_c_*) to transition to a processing state, during which they neither forage nor disperse. Each processing individual eventually reproduces (at rate 1*/h_c_*) giving rise to two foraging consumers. Both foraging and processing states experience identical intrinsic mortality rates (*d_c_*).

Since we want to isolate the effect of the metapopulation size, we do not allow the ‘disappearance’ of patches driven by the local extinction of the sedentary resource (which makes a patch unsuitable for consumers and thus lost from the metapopulation for all intents and purposes). We, therefore, allow a weak external influx of resource (at rate *E_r_*), which ensures eventual re-population of patches where the resource species goes extinct.

The six ecological parameters (*ω_r_, K_r_, E_r_, ω_c_, h_c_, d_c_*) are exogenously fixed across the metapopulation and do not change in the course of evolution. However, their values vary across different evolutionary simulations and serve as control parameters for ecology sampling.

### Between-patch dispersal dynamics

At each simulation step, foraging consumers independently choose between continuing to forage versus dispersing, based on their dispersal strategy and the available information about the local patch state. Thus, the decision-making timescale is the fastest timescale in our model, followed by the intermediate timescales of patch residence, demography, and resource depletion/recovery; the evolutionary timescale is the slowest.

Dispersing comes at the cost of forgoing foraging for one simulation step (the dispersing step). We examine three sensory models:

- *Uninformed*: no information can be gathered so that, if dispersal evolves, it occurs at a constant rate.
- *Conspecifics sensing*: consumers sense the total number of consumers in the patch.
- *Resource sensing*: consumers sense the amount of resource in the patch.

The uninformed model uses a state-independent dispersal rate *m*_0_. Both informed models (conspecifics and resource sensing) employ a sigmoid dispersal rate *m*(*I*), see Fig. 1C.

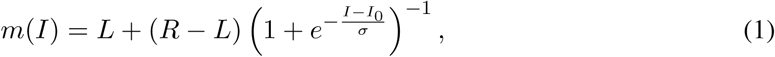

where, depending on the model, *I* represents either the number of consumers (*N_c_*, for conspecifics-sensing) or the amount of resource (*N_r_*, for resource-sensing) in the patch, *L* and *R* are the limits of the dispersal rate for very low and very high values of sensory input, respectively, *I*_0_ is the position of the inflection point, and *σ* is the width of transition.

### Evolution

Dispersal strategies are defined by either one (*m*_0_; uninformed model) or four (*L, R, I*_0_*, σ*; informed models) parameters. All parameters are subject to evolution, which we simulated via an iterative mutation-selection process: more successful strategies leave more offspring than their competitors, while their mutants further explore the parameter space in subsequent selection rounds. Over evolutionary time, this process yields strategies that optimally use sensory information given the ecological conditions. Because the sigmoid function can take various shapes depending on its parameters (constant, near-linear, step-like, etc.), evolution acting on those parameters can capture a diversity of dispersal behaviors.

### Classification of dispersal strategies

We characterize the dispersal strategies (*m*(*N_r_*) or *m*(*N_c_*)) evolved at diverse ecological conditions by testing which of their features are necessary for their evolutionary success under the corresponding combination of ecological parameters. To this end, we competed each strategy against “knockouts” that were modified to isolate specific features (Fig. 1C). To do this classification systematically, we ask the following questions:

1. Is dispersal beneficial in this ecology? If it is, the evolved strategy should out-compete the “knockout” with *No-dispersal* (*m*(*I*) = 0, gray lines on Fig. 1C). If it does not, then the evolved strategy is a “*No-dispersal*” strategy.
2. Is it beneficial for the dispersal decision to account for the patch state? If it is, the evolved strategy should out-compete “knockouts” that are insensitive to information (*m*(*I*) = *const*, brown lines on Fig. 1C). If it does not, then we call the evolved strategy “*Ignorant*”.
3. Is it always beneficial to stay in a high quality patch? If true, the evolved strategy should *not* outcompete the “knockout” with no base-line dispersal (min(*m*(*I*)) = 0, pink lines on Fig. 1C). If it does, we call such a strategy “*Anxious*”.
4. Is it always beneficial to leave a low-quality patch? If true, the evolved strategy should *not* out-compete the “knockout” with no saturation of the dispersal rate at low quality conditions (*m*^′′^(*I*) *>* 0, turquoise lines on Fig. 1C). If it does, we call such a dispersal strategy “*Saturated*”. If the evolved strategy does not outcompete the “knockout”, then it has an “*Unsaturated*” dispersal behavior, which recovers the marginal value theorem (MVT) prediction.

A more detailed model description can be found in Appendix 1, and the formal definitions of the “knockout” strategies are provided in Appendix 2.

## 3 Results

For all three models, we studied the evolution of dispersal strategies in more than 13000 combinations of the ecological parameters, which we refer to henceforth as ‘ecologies’ (see ecology sampling details in Appendix 1). To establish a baseline expectation for no dispersal, we also studied the ecological dynamics in an isolated patch under the same ecologies. Many of these ecologies were not viable. By training a decision-tree classifier we found that, quite surprisingly, the viability of the metapopulation in a given ecology can be inferred with a high accuracy (F1-score *>* 0.8) from just two features of the consumer-resource dynamics in an isolated patch, in the same ecology: the equilibrium consumer population, 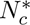, and the equilibrium fraction of the total available resource, 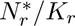 ; see Appendix 3 for a detailed analysis. Taking advantage of this insight, we visualized the multi-dimensional space of ecologies by projecting it to a 2-dimensional plane, see Fig. 2B,C,F-H, which allowed us to extract comparative insights across models.

**Figure 2:**
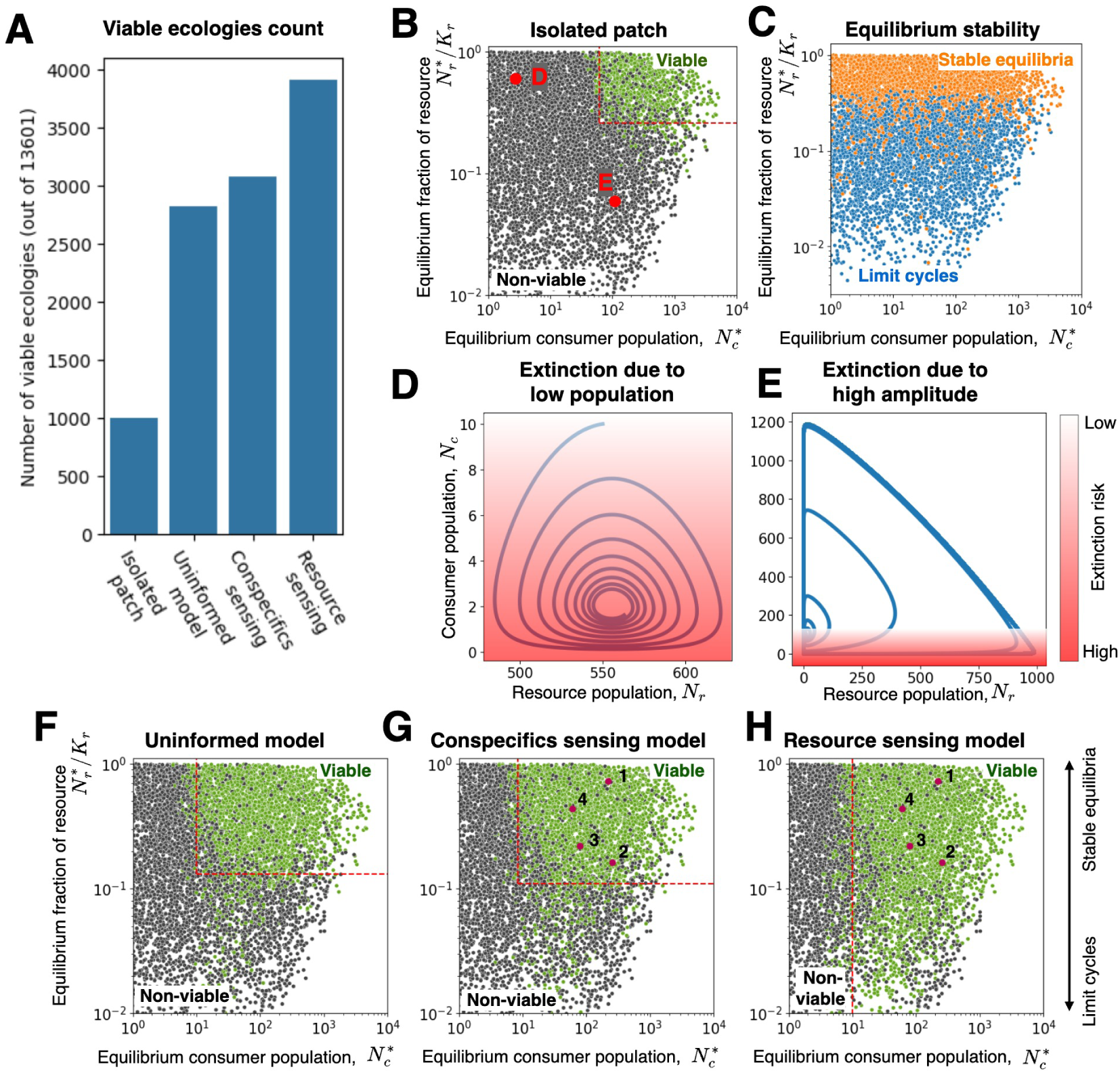
Informed dispersal improves the viability of trophic meta-populations. **A** Number of sampled ecologies capable to support a viable consumer population in different models. **B** Map of ecologies with viable (green) and non-viable (black) consumer populations in an isolated patch. Red dots are locations of ecologies shown on panels B and C. **C** Map of equilibrium stability of an equivalent ODE model, see Appendix 1, section 3. **D,E** Two ways how a population may go extinct in a patch: due to low equilibrium population size (D) or due to a limit cycle with high amplitude (E). Blue lines are the population trajectories in the equivalent ODE model. The intensity of red shade qualitatively indicates the extinction risk in a stochastic dynamics model. **F - H** Maps of ecologies with (non-)viable meta-communities in each of considered sensory models. Dashed lines show the rectangular region of viable ecologies, according to the decision tree classifier described in Appendix 3. Points 1-4 indicate four representative ecologies considered throughout the main text. Parameters used: *M* = 20, *d_c_* = 1.0, *E_r_* = 0.1, while *ω_c_, h_c_, ω_r_, K_r_* vary among samples and *m*_0_*, L, R, σ, I*_0_ use evolved values. are provided in Appendix 3). Survival under uninformed (Fig. 2F) and conspecifics-sensing (Fig. 2G) models are subjected to similar restrictions, while resource-sensing uniquely eliminates the resource fraction constraint (Fig. 2H).

All results below are obtained for a metapopulation with *M* = 20 patches, but we confirmed the robustness of our findings for both smaller (*M* = 2) and larger (*M* = 50) meta-communities (see Appendix 4).

### 3.1 Steady state consumer and resource levels predict consumer viability in an isolated patch

We first start by analyzing an isolated patch, where in an infinite population ODE the only two possible outcomes are stable equilibria or limit cycles (Fig. 2C). In the finite population simulations, as expected, we found extinction either due to demographic stochasticity at small consumer populations (Fig. 2B,D), or due to limit cycles with large enough amplitude (Fig. 2B,E). An unexpected result was the predictive power of the resource level for the extinction of consumers in ecologies exhibiting such analytically intractable limit cycles (Fig. 2B,C).

### 3.2 Dispersal improves consumer viability; informed dispersal under resource sensing maximizes it

Comparing across models, we see that the possibility of dispersal provided by a meta-population setting extends the range of viable ecologies relative to an isolated patch, and that informed dispersal extends it more than uninformed, with the resource-sensing model being the most efficient in that regard; see also Fig. 2A for a quantification. Specifically, being viable in an isolated patch (Fig. 2B) requires a larger equilibrium consumer population and a larger fraction of the resource at equilibrium than survival in a meta-population with any sensory model (Fig. 2F-H; inferred viability thresholds

The number of patches in the metapopulation affects viability (i.e., increasing the number of patches increases the number of viable ecologies), but it does not change the evolution outcomes, see Appendix 4.

### 3.3 Under uninformed dispersal, abstaining from dispersal could be an optimal strategy

Optimal dispersal rates (*m*_0_) correlate with the consumer population size and the equilibrium stability. We find elevated dispersal rates to evolve at ecologies with either small consumer populations (low 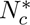) or unstable equilibrium (low 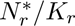), see Fig. S6 in Appendix 5. These findings match the patterns observed in [Pillai et al., 2012, Laroche et al., 2016], where the optimal dispersal rate correlates with the extinction risks.

In ecologies with large populations at a stable equilibrium (high 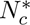 and 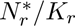) where these risks are negligible, we find evolved dispersal rate (*m*_0_) to be so low that the optimal strategy is unable to outcompete the sedentary strategy with *m*_0_ = 0. These strategies form the *No-dispersal* class, see Fig. S7 in Appendix 5. In these ecologies, consumers experience neither periods of starvation, which would promote escaping from the current patch, nor periods of overabundance, which would provide an opportunity for an invasion into other patches. At any time, moving to another patch is not expected to bring a strong enough improvement in surrounding conditions to warrant paying the cost of dispersal – skipping a round of foraging. As a result, dispersal is disadvantageous, and the *No-dispersal* strategies evolve. The existence of *No-dispersal* class and its ecological prerequisites are not unique to the uninformed dispersal but instead are shared across all three sensing models, see Fig. 3A,C and Fig. 4A,G.

**Figure 3:**
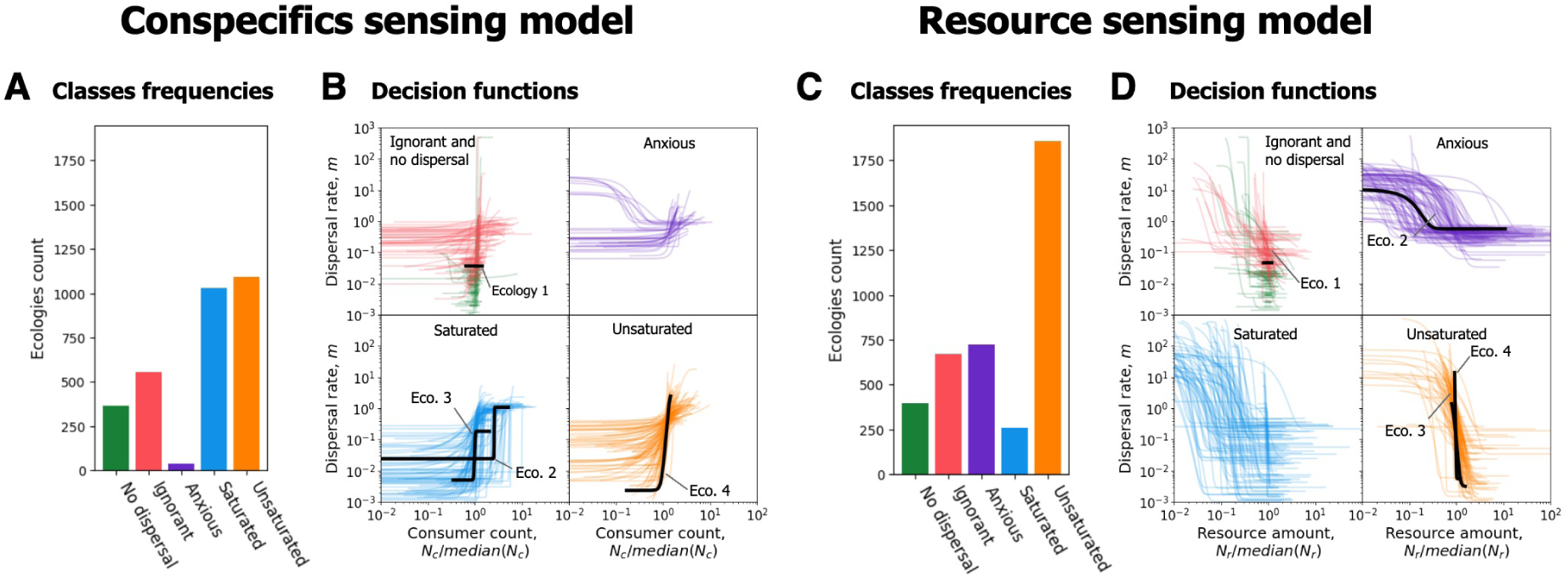
Evolved decision functions group into five classes: *No-dispersal*, *Ignorant*, *Anxious*, *Saturated*, and *Unsaturated*. **A,C**: frequency of each class in the conspecifics and resource sensing models. **B,D**: decision functions in each class in two models. Colored curves show the sample of 100 decision functions (*m*(*I*)), which belong to the given class. The support of each decision function indicates the range of argument values actually experienced by the population in a simulation. Solid black curves are the decision functions evolved in four representative ecologies, which demonstrate the most prevalent combinations of strategy classes that arise across the two informed models. Parameters used: *M* = 20, *d_c_* = 1.0, *E_r_* = 0.1, while *ω_c_, h_c_, ω_r_, K_r_*vary among ecologies and *L, R, σ, I*_0_ use evolved values. Parameters of the representative ecologies can be found in Appendix 6.

**Figure 4:**
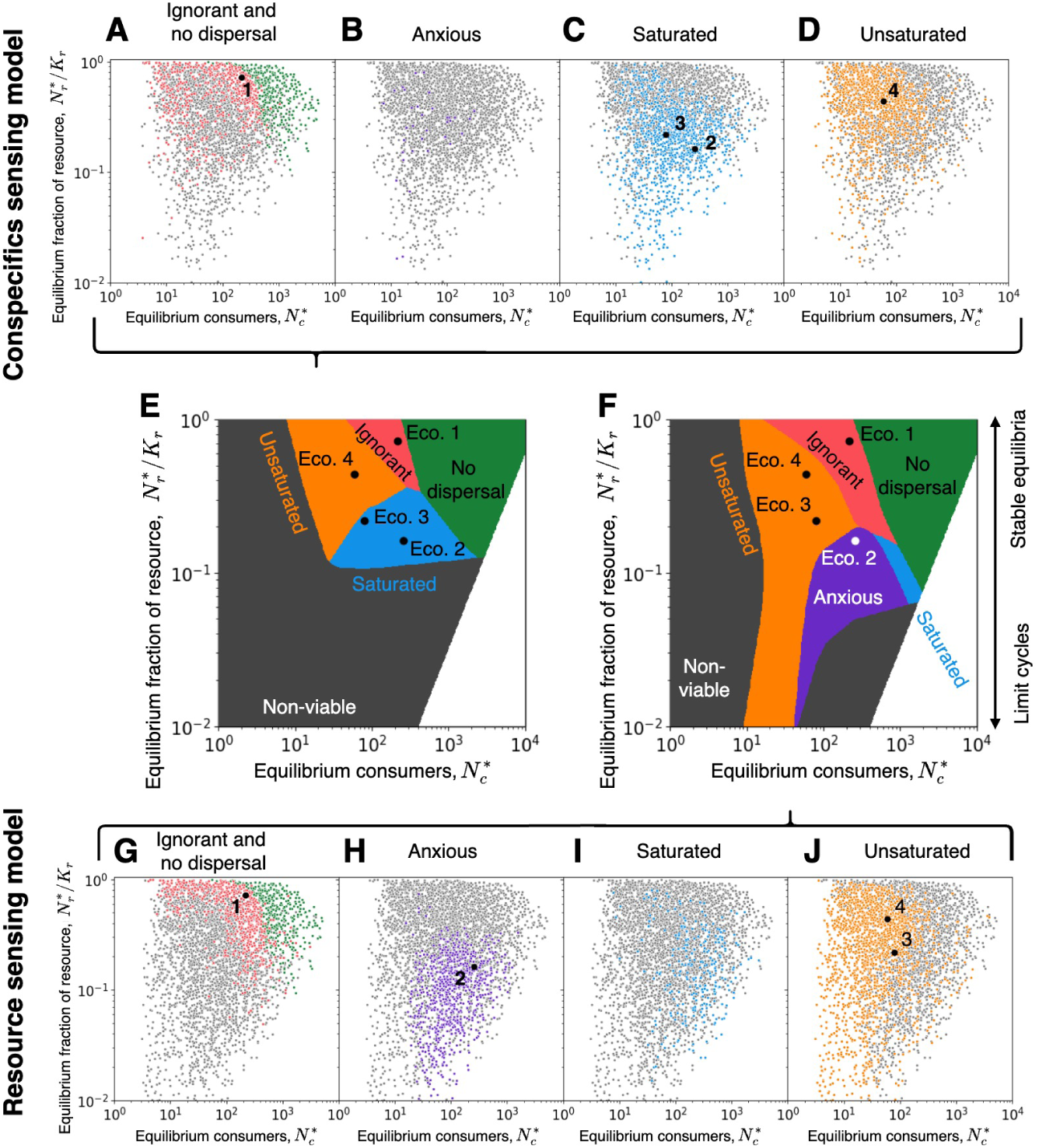
Each classes of dispersal strategies evolves in a distinct region of the ecology space. **A-D** Locations of all viable ecologies promoting a dispersal strategy of a given class in the ecology space under the conspecifics sensing model. Colored dots - ecologies promoting one of five classes of dispersal strategies, grey dots - all other viable ecologies. Black circles - locations of four example ecologies shown together with the class they belong to. All panels show a projection of the 4-dimensional ecology space to the plane: equilibrium consumers 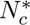 vs equilibrium fraction of resource 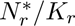. **G-J** The same under the resource sensing model. **E, F** Regions of optimality of each class in conspecifics sensing model (E) and resource sensing model (F) as interpolated by a neural network classifier. Parameters used: *M* = 20, *d_c_* = 1.0, *E_r_* = 0.1, while *ω_c_, h_c_, ω_r_, K_r_*vary among ecologies and *L, R, σ, I*_0_ use evolved values.

### 3.4 Under informed dispersal, evolved strategies group into five qualitatively distinct classes: *No-dispersal*, *Ignorant*, *Anxious*, *Saturated*, and *Unsaturated*

We classify the dispersal strategies that evolve across our simulations in the two sensing models into five groups (Fig. 1D), based on the results of their competition against systematically designed “knock-outs” (Fig. 1C, and Appendix 6). The strategies belonging to the first, *No-dispersal*, class cannot outcompete a “knockout” that does not disperse; consequently, any dispersal they might exhibit is not adaptively significant. The strategies belonging to the *Ignorant* class cannot outcompete a “knockout” that is insensitive to information; consequently, for *Ignorant* strategies, any dependence of the dispersal rate on the patch state does not yield a selective advantage. The remaining three classes evolve to use the available information. Strategies belonging to the *Anxious* class derive their success from the decision to disperse from a high-quality patch (resource-rich or underpopulated). Strategies belonging to the *Saturated* class derive their success from their bounded (i.e., capped) dispersal from low quality patches. And, finally, strategies belonging to the *Unsaturated* class, derive their evolutionary success from gradual acceleration of the departure rate with decreasing patch quality; this class, therefore, contains, among others, the strategy predicted by the MVT. Henceforth, we focus only on the four most common classes in each sensing model: under conspecifics-sensing we ignore the *Anxious* strategies due to their low numbers; under resource-sensing, we ignore the *Saturated* strategies due to their low numbers and because they represent a transitional state between *Anxious* and *Ignorant* strategies rather than being a distinct group.

The shapes of the evolved decision functions reflect the class definitions (Fig. 3B,D). Both *Ignorant* and *No-dispersal* decision functions are predominantly flat, with the latter exhibiting minimal dispersal rates. *Anxious* strategies consistently exhibit substantial dispersal from good-quality patches (with, expectedly, even higher dispersal from low-quality ones). *Saturated* decision functions limit dispersal from low-quality patches. *Unsaturated* functions vary in shape but, relative to the other classes, they are the most sensitive to deviation from median patch conditions: i.e., the values of the inflection point (*I*_0_) evolve to be close to the median state of a patch, calculated across time in the stochastic simulation of the meta-population. The meaning of this emergent feature will crystallize only in the next section, when we look at the ecological conditions that select for *Unsaturated* strategies.

### 3.5 Which class of strategies is evolutionarily optimal is determined by the ecological conditions

The five classes not only exhibit characteristic decision functions, but also occupy distinct regions of ecological parameter space. Our 2D projection of that space showing the viability of ecologies (Fig. 2) also reveals the ecological conditions favoring each strategy class (Fig. 4). We see that *Ignorant* and *No-dispersal* strategies evolve in ecologies where the population in an isolated patch would exhibit stable equilibria, whereas *Saturated* and *Anxious* strategies evolve when the population would exhibit limit cycles. Interestingly, *Unsaturated* strategies can evolve in either of these broad regimes.

In ecologies supporting large, stable consumer populations (high 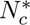 and 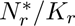), *No-dispersal* strategies evolve, similar to uninformed model, see Fig. 4A,G. There, the minimal benefits of dispersal cannot outweight its cost (skipped foraging time), thus the ability to sense the local environment is of a little use.

In ecologies supporting somewhat smaller, yet stable consumer populations (intermediate 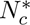, high 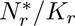), a lucky migrant may occasionally fixate in the destination patch; eventually this process can lead to the complete displacement of the sedentary, *No-dispersal* strategy. In these ecologies, *Ignorant* strategies emerge, see Fig. 4A,G, which gain their selective advantage not from a clever use of sensed information, but through stochastic fixation of randomly departed migrants.

In ecologies supporting small, stable consumer populations (low 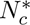 and high 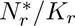) the magnitude of the stochastic fluctuations becomes so high that dispersing at a constant dispersal rate is more likely to result in (local) extinction than sensing the local information and making a dispersal decision based on it. Since in these ecologies correlations across patches remain weak (Fig. 5A,B), dispersal results in arrival at an average quality patch. *Unsaturated* strategies, with their evolved sensitivity to median patch conditions, are precisely the kind of strategies that evolved to make a dispersal decision based on how the current patch compares to the average destination, which is why we find this region of parameter space (low 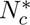, high 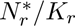) overwhelmingly dominated by *Unsaturated* strategies in\ both sensory models (see Fig. 4E,F, and Fig. 5C). In particular, the *Unsaturated* class contains the categorical decision strategy—stay as long as the local patch is better than average and instantaneously leave as soon as the average destination becomes better. In the resource sensing model, this is exactly the MVT strategy.

**Figure 5:**
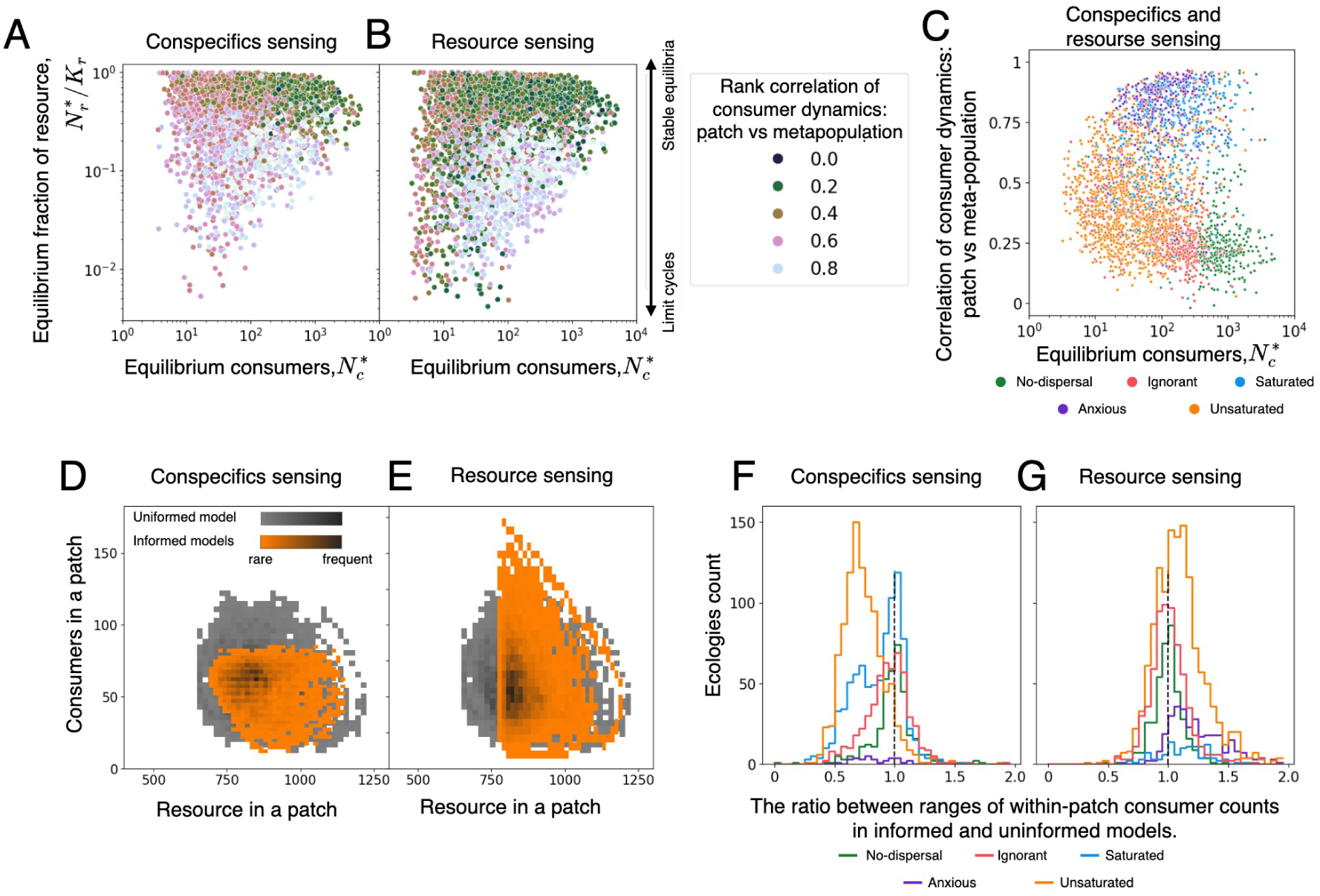
Classes of dispersal strategies are determined by spatial correlations and the patch population size. Panels **A, B**: maps of correlations between patch and meta-population consumer dynamics in ecological space for conspecifics (panel A) and resource sensing models (panel B). Panel **C**: joint map of all classes in both sensing models. Each point represents an ecology and its color shows the class of the evolved dispersal strategy. Panels **D, E**: heatmaps of the numbers of resource and consumers in a patch in the course of population dynamics in representative ecology 4. Darker color indicate more frequent patch state. Gray shades - uninformed model, orange shades - informed model: panel D for conspecifics sensing, panel C for resource sensing. **F, G**: Distribution of the ratio between ranges of within-patch consumer counts over the course of population dynamics in informed and uninformed models across ecologies separated by class. Values larger than 1 indicate amplification of resource count range in the informed model, compared to uninformed; values below 1 indicate dampening of variation. Vertical dashed line indicate value equal to 1. Only ecologies with viable population in both considered models are taken into account. Parameters used: *M* = 20, *d_c_* = 1.0, *E_r_* = 0.1, while *ω_c_, h_c_, ω_r_, K_r_* vary among ecologies and *L, R, σ, I*_0_ use evolved values.

In ecologies exhibiting limit cycles (low 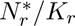), we find the optimal dispersal strategies to be shaped by emergent dynamics at the meta-population scale. Here, the first patch in the metapopulation that reaches the peak of its population size is capable of sending out multiple migrants (and typically does so); the elevated emigration from that patch populates lagging patches and pushes them towards their own peaks. This patch-synchronizing effect is countered by within-patch stochastic fluctuations, which occur independently in each patch and are, therefore, de-synchronizing. At low consumer populations, 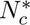, migration flows are weak while stochastic fluctuations are relatively strong and hence, they overcome the synchronizing effect, which means that dispersal from a resource-poor/overpopulated patch results in arrival to an average-quality one. Consequently, *Unsaturated*strategies also dominate this region of parameter space (low 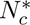, low 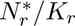) in both sensory models (see Fig. 4E,F, and Fig. 5C).

When the synchronizing effect overcomes the within-patch stochastic fluctuations (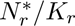 are low but 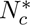 are not), patches will go through peaks and valleys of consumer and resource abundance at similar times, i.e., consumer dynamics are highly corrleated across patches (Fig. 5A,B). Here, *Unsaturated* strategies are at a disadvantage because the expected state at the destination patch becomes time-dependent and, therefore, cannot be encapsulated by one value (the average patch quality). Moreover, a strongly correlated patch dynamics means that destination patches are in a similar state to the origin patch. Nonetheless, the correlation is not 1, so there will sometimes be patches that are lagging in their dynamics and could be suitable for consumers; hence, there is still advantage to dispersing from low quality patches. How high the dispersal should be from a low-quality patch depends on the ability of the model to predict the future conditions in such a patch based on its current state, which is why we find different strategies evolving in the conspecifics-sensing (*Saturated* strategies) versus resource-sensing models (*Anxious* strategies).

Under conspecifics sensing, a resident sensing an overpopulated patch will not be able to infer whether that patch is overpopulated and declining or overpopulated but still growing (i.e., yet to reach its peak, which exact magnitude is stochastic and thus unpredictable). Therefore, under conspecifics sensing, consumers will tune their dispersal rate to balance the possible benefits of leaving a declining patch with the costs of leaving too early from a growing patch. This balancing act results in the emergence of *Saturated* strategies with a minimal dispersal from underpopulated patches and capped dispersal from overpopulated patches.

Under resource sensing, a resident sensing low resources will correctly infer that the patch consumer population is in decline. Therefore, there is no balancing required and dispersal can be very high or even uncapped. *Anxious* strategies emerging there in ecologies with limit cycles exploit the correlated meta-population dynamics. A resident sensing a resource-rich (hence, growing) patch may expect other patches to be growing as well, thus the risks of dispersal from a resource rich patch are much less than in other models. At the same time, an opportunity to move from a growing but overpopulated patch, approaching the peak of its population and inevitable collapse, to a patch lagging behind on the limit cycle does not only boost the immediate growth rate (lagging patch has more resources) but also allows entering the decline phase later than a resident, which stays until the growth pahse ends.

### 3.6 Informed dispersal impacts within-patch dynamics differently relative to uninformed dispersal

To determine whether the ability to use information has any impact on within-patch dynamics, we compare the dynamics emerging in a given ecology under different sensing models. The largest difference is, again, in the viability of consumer population: many ecologies viable under the resource sensing model are non-viable in two other models (cf.Fig. 2F-M). We return to this phenomenon towards the end of this section but for now we select for our analysis only the ecologies, which are simultaneously viable under the uninformed and either of informed models.

The classes of *No-dispersal* and *Ignorant* strategies are comprised of, effectively, uninformed strategies and, therefore, produce similar dynamics across all three models, see Appendix 7 for the detailed view on representative ecology 1 and Appendix 8 for the summary among multiple ecologies. Furthermore, for all five classes, the long-term averages of the consumer and resource population sizes, measured over time intervals longer than the period of limit cycle, differ insignificantly between informed and uninformed models, see Appendix 8. In a closed system, dispersal cannot change the scale of population size determined by the environment and the ecological interactions. However, we find the magnitude of variation around these averages to be systematically different between informed and uninformed models, with each sensing model exhibiting its own character of changes.

Under the conspecifics sensing model, in *Unsaturated* class we find the population fluctuations to be dampened compared to the uninformed model, (see Fig. 5D for an example provided by the representative ecology 4 and Fig. 5F for the summary among multiple ecologies). Here, elevated dispersal from an overpopulated patch reduces the local consumer density, which, in turn, immediately decreases the dispersal rate. As a result, *Unsaturated* conspecifics-sensing strategies suppress fluctuations of the consumer population allowing both the consumer and the resource populations to hover closer to the equilibrium state than would be possible under the uninformed model.

The impact of the *Saturated* class (conspecifics sensing) is bi-modal: some ecologies suppress variation in numbers of both resource and consumers, while others keep them at the levels similar to the uninformed model, see Fig. 5F. The source of this bi-modality is that the observed variation in the population size comes from two sources: the limit cycle dynamics and the stochastic fluctuations around the limit cycle trajectory. In ecologies, where variation in population size is mainly due to fluctuations around a limit cycle (i.e. with smaller population size), these fluctuations can be suppressed by the informed dispersal, as it happens in the *Unsaturated* class above. By contrast, in the ecologies with large population, the variation in its size is mostly driven by the amplitude of the limit cycle itself, with fluctuations playing a minor role. Given the high correlation among patches, any level of dispersal from an over-populated patch will be compensated by the similar level of immigration from other patches at similar states. As a result, two subgroups of the *Saturated* class correlate with the consumer population size (see Fig. S12 in Appendix 8).

The impact of the dispersal with resource sensing is more nuanced. For instance, the *Unsaturated* class amplifies the variation in consumers numbers but suppresses the variation in resource amount, as demonstrated by the representative ecology 4 on Fig. 5E and summarized for multiple ecologies in Fig. 5G and Fig. S11 in Appendix 8. Here, the resource-rich patches experience an imbalanced flow of migrants: the negligibly low outbound migration cannot compensate the incoming migration flow from resource-poor patches. Thus, the resource-rich patches accumulate larger population of consumers than under the uninformed model resulting in a larger variation of consumer numbers. At the same time, the elevated dispersal from resource-poor patch prevents the resource level from falling too low. Naturally, this improves viability of ecologies exhibiting limit cycles under the resource-sensing model, see Fig. 2H. The high dispersal level from resource-poor patches means that only a few consumers will be foraging when the resource amount is low. Reduced foraging pressure means that the local extinction of the resource population is less likely and its regrowth proceeds faster, which in turn keeps the consumer population viable in the long-term.

In the *Anxious* class, the dynamics is similar – consumer variation is amplified because resource-rich patches receive more migrants than they send out, see Fig. 5G. Compared with *Unsaturated* class, very high dispersal rates from resource poor patches increase the amplification effect (more migrants from each resource poor patch) but base-line dispersal and high correlation among patches reduce its strength (permanent outflow of migrants and shorter period with migration flow imbalance) – overall, the strength of the amplification remains the same.

## 4 Discussion

In this work, we establish a broad baseline for the evolution of informed foraging dispersal. To this end, we consider a mechanistically minimal model of a trophic meta-community, where the evolution of flexible dispersal strategies is driven by the ecological consequences of dispersal decisions. To determine the breadth of possible strategies, establish their robustness, and meaningfully compare them, we run the eco-evolutionary process over a large number of possible configurations of the resource-consumer ecological dynamics. As a result, we uncovered a rich space of dispersal strategies. Among them, we recovered the dispersal dynamics predicted by classical theory (MVT) in the *Unsaturated* class. In addition, our model also uncovered the existence of four other classes of strategies: *No-dispersal*, *Ignorant* (i.e. a strategy evolved to use no available information, thus recapitulating the evolution of uninformed strategies), *Saturated* (i.e. exhibiting delayed dispersal from unfavorable conditions), and *Anxious* (i.e. exhibiting some non-zero dispersal from a favorable patch). Each of these five classes is characterized by a distinct shape of the decision function (Fig. 3) and evolves under specific ecological conditions (Fig. 4).

The diverse strategies we find in this work have been previously found empirically. The *Ignorant* dispersal has been reported for foraging of apes [DiGiorgio et al., 2020, O’Bryan et al., 2020], teals [Tamisier, 1974], and insects movement [Sappington, 2024]. The limited dispersal from overpopulated patches, exhibited by *Saturated* strategies, is reported for small rodents [Barash, 1974, Fairbairn, 1978, Krebs et al., 1978, Slade and Balph, 1974, Michener and Michener, 1977, Gaines and McClenaghan, 1980] and bluegill [Crowley et al., 1990]. However, they were previously reported as unrelated and isolated violations of the MVT predictions. Our results show that these findings are not just deviations from the single theoretically optimal MVT strategy but represent alternative classes of potentially optimal dispersal strategies emerging in the course of the eco-evolutionary dynamics under different set of ecological parameters.

This suite of non-MVT strategies have also been previously reported in the theoretical literature, but not in a meaningful comparison with each other. The *Ignorant* strategies were predicted to evolve in species with strong inter-generational effects [McNamara and Dall, 2011]. A *Saturated* strategy has been suggested to be the true optimal strategy in single-species meta-populations with externally driven patch quality [Hovestadt et al., 2010]. Unwillingness to settle in the resource-rich conditions, exhibited by *Anxious* strategies, has been theoretically predicted to evolve in meta-communities with high temporal variability of the patch populations [Khattar et al., 2024]. However, without an encompassing baseline in which to compare and evaluate all such dispersal strategies, it has been challenging to put them into perspective. An attempt to provide a comparison was performed in [Khattar et al., 2024], but only in a setting with externally driven patch quality and exponential decision functions.

Our framework provides a complementary comparative perspective, with the ecological and evolutionary dynamics being endogenous and the decision-functions more flexible. Moreover, we perform an extensive parameter space exploration by sampling two orders of magnitude more ecological conditions. Such a broad setting recovers five classes of strategies and allows us to extract a meaningful comparison (Table 1). For instance, which strategies evolve depends on the magnitudes of temporal and spatial demographic variations relative to the equilibrium state. Low temporal variation (weak fluctuations around stable equilibrium) promotes the *Ignorant* and the *No-dispersal* strategies. High temporal variation combined with a low spatial variation (correlated limit cycles) lead to either *Saturated* or *Anxious* strategies, depending on the available information. Finally, high temporal and spatial variations (uncorrelated limit cycles or large fluctuations around stable equilibrium) give rise to *Unsaturated* strategies.

**Table 1:**
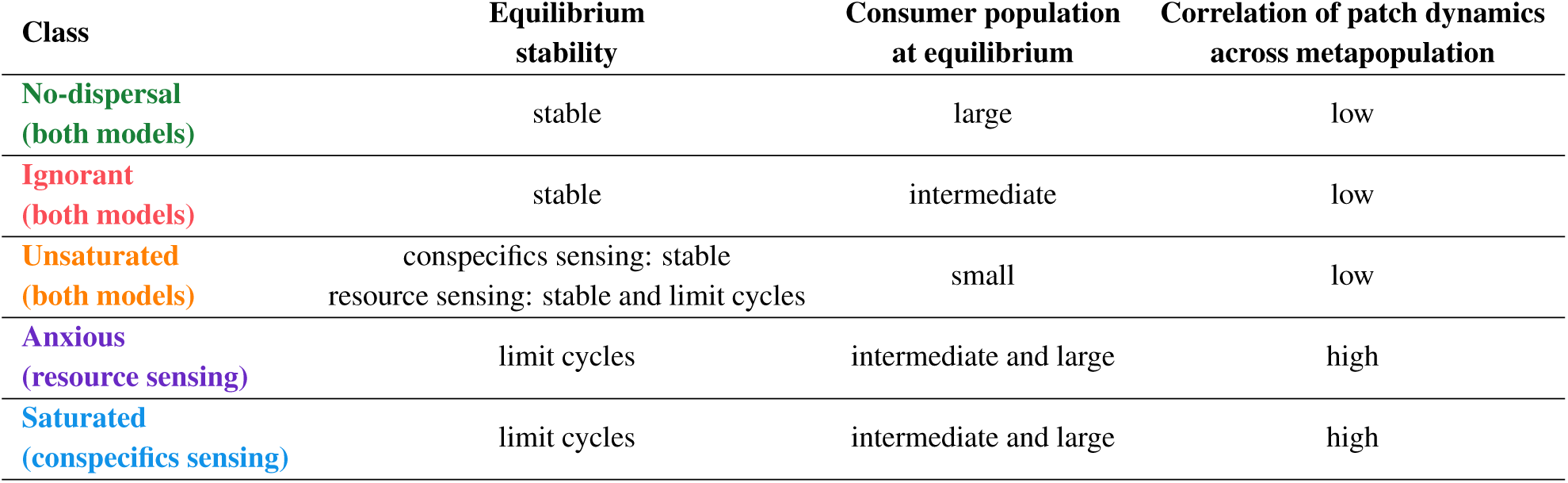
Summary of ecological conditions promoting different classes of dispersal strategies.

| Class | Equilibrium stability | Consumer population at equilibrium | Correlation of patch dynamics across metapopulation |
| --- | --- | --- | --- |
| **No-dispersal (both models)** | stable | large | low |
| **Ignorant (both models)** | stable | intermediate | low |
| **Unsaturated (both models)** | conspecifics sensing: stable<br>resource sensing: stable and limit cycles | small | low |
| **Anxious (resource sensing)** | limit cycles | intermediate and large | high |
| **Saturated (conspecifics sensing)** | limit cycles | intermediate and large | high |

This type of insight also allows us to understand which strategies cannot evolve. Although every conceivable sigmoid function is theoretically permissible within our model, the strategies that actually evolve represent only a subset of these possibilities. For instance, strategies with negative sensitivity — where better local conditions lead to more active dispersal — are virtually absent from our results. These strategies, although they may sound maladaptive, were in fact predicted to evolve in large populations undergoing an externally driven non-stationary dynamics without any spatial correlation among patches [Parvinen et al., 2023]. By contrast, endogenously driven population dynamics prevents the simultaneous occurrence of these three conditions (large population, non-stationary dynamics, and no spatial correlations) because large non-stationary populations evolve strategies that enhance spatial correlations. This finding, therefore, clarifies that the presence of a strong external driver is, in fact, a necessary condition for negative sensitivity strategies to emerge. Although here we took the complementary lens of an entirely endogenous process, our framework can be easily extended to encompass externally-driven conditions, by, e.g., allowing spikes of exogenous resource immigration and allow even broader comparisons of evolved strategies.

Our work further sheds light on the effect of different types of sensed information, since for each ecology we probe the evolution of dispersal for three different sensory models: specifically, consumers could be sensing the local amount of resource, the local density of conspecifics, or neither (i.e. uninformed). The type of perceived information determines the viability of ecology (see Fig. 2). Sensing the amount of conspecifics contributes to the survival only a little compared with uninformed dispersal, while sensing the amount of resource allows population to thrive in otherwise non-viable ecologies (see Fig. 2F-H and Appendix 3). There, the conspecifics sensing allows to assess the current state of the local population, while the resource sensing effectively assesses the rate of its change (growth or decline of the local population) – the latter is more useful in anticipating the immediate future. The available information also influences the decision function - the way, how the information should be turned to action. In ecologies with inherently non-stationary dynamics - the limit cycles, different sensory models bring forward different strategies. There, optimal decision functions belong to *Saturated* class in the conspecifics sensing model but *Unsaturated* or *Anxious* in the resource sensing model. At the same time, ecologies with large and stable populations promote *Ignorant* and *No-dispersal* classes in all sensory models. There, the information provided by sensors are simply not useful for a dispersal decisions, so all three sensory models align in their predictions. Of course, some species may be able to use multiple sources of information. It would be an interesting future direction to explore how individuals would evolve to weigh multiple types of information (e.g., in our context, conspecifics versus resource sensing) and what types of dispersal strategies would emerge in that context.

In this work, we separated the timescales of decision making, demographic, and evolutionary processes. The fastest is the timescale of individual decision-making, which takes only a single timestep. The demographic timescale is slower: reproduction rates vary across ecologies, but in all cases an average consumer individual makes 10 dispersal decisions during its lifetime. The mutation-selection timescale is slower still, at roughly 500 average consumer lifetimes. Such a long selection round allows capturing the impact of emergent population dynamics, such as synchronization of limit cycles across the meta-population. Finally, the evolutionary timescale takes about 50000 average lifetimes (100 times slower than mutation/selection), which is necessary for random mutations and directed, but noisy, selection to be able to approximate the optimal strategy. This timescales separation reflects the intuitive hierarchy of individual actions, population dynamics, and evolution, and makes it possible to disentangle the interplay of interactions between different processes constituting the eco-evolutionary feedback loop. However, it is important to note that in natural populations the timescales are not always separated. For instance, decision-making and demographic timescales overlap in annual plants, which make only one decision about their dispersal of their seeds in the course of their lifetime [Howe and Smallwood, 1982]. Mutational and demographic timescales overlap in bacterial strains with hy-permutation, in which Petri dish colonies grown from a single cell are notably heterogeneous [Barnett et al., 2025]. Such cases deserve their own separate theoretical treatment.

Our results offer a revised perspective on the empirical probing of dispersal. Since decision functions are difficult to accurately measure in experiments, proxy measures are used instead. Following the predictions of MVT, these measures typically search for the threshold of leaving; for instance, giving-up-density [Brown, 1988, Bedoya-Perez et al., 2013] measures the minimal amount of available resources justifying staying in the current patch. However, this characteristic MVT threshold for leaving the patch at the marginal gains level exists only in the *Unsaturated* class of dispersal strategies, see Fig. 3B,D. For *Saturated* strategies, the dispersal threshold is not correlated to the marginal gains, resulting in overstaying behavior under the MVT interpretation. The threshold featured by *Anxious* strategies does not separate staying from departing, and there are no thresholds for *Ignorant* and *No-dispersal* strategies at all. As a complementary approach to giving-up-densities and other threshold-focused proxy measures, we suggest that additional tests might help distinguish among the different classes. First, a comparison of dispersal behavior from a low- versus a high-quality patch allows helps narrow down possibilities. If dispersal proceeds in the same way in both cases, it is either an *Ignorant* or a *No-dispersal* strategy. If dispersal behavior differs between low- and high- quality patches and individuals leave a high-quality patch, then the strategy is *Anxious*; otherwise it belongs to either the *Saturated* or the *Unsaturated* classes. Next, to distinguish between the last two classes, an additional test is needed to probe whether dispersal behavior is sensitive to a patch state at low-quality conditions: *Saturated* strategies are insensitive to patch quality once it falls low enough, but *Unsaturated* strategies are not. This can be done, for instance, via an additional comparison of dispersal behavior at low quality conditions of a different magnitude, e.g. at the complete absence of resource. If the experimental setup does not allow the inference of dispersal rates with sufficient accuracy, a proxy measure such as residence time could be used instead. This shift away from threshold-centric proxies provides a framework to disentangle the diversity of dispersal strategies in nature.

We are grateful to members of Tarnita lab at Princeton and Theoretical Biology Department at Max Planck Institute for Evolutionary Biology, as well as to Simon Levin for constructive feedback. The detailed description of the models and analysis methods are presented in Appendices 1–8.

