## Appendices for "Evolution of informed dispersal strategies in trophic meta-communities"

### Supplementary information to the manuscript: Evolution of informed dispersal strategies in trophic meta-communities

**2 Knockout competition** **13**

**3 Viability of consumer populations across ecologies and sensory models** **15**

**4 The same kinds of strategies evolve in metapopulations of different sizes** **18**

**5 Evolution of dispersal rate in the uninformed model** **20**

**6 Classification of the decision functions** **22**

**7 Resource-consumer dynamics in representative ecologies** **24**

**8 Comparison of within-patch dynamics between informed and uninformed sensing models** **26**

#### 1 Complete model description

We consider a meta-community inhabiting $M = 20$ identical patches forming a fully connected graph. These patches are populated by two species: the resource and the consumer. Resource individuals are all identical and immobile. Consumer individuals can move between patches and may differ in their dispersal strategies, see section 1.1. Population dynamics within each patch are governed by trophic interactions, see section 1.2 for the model implementation and section 1.3 for the analytical approximation. We implement mutation-selection process to find the most advantageous dispersal strategies of the consumer species, see section 1.4. The character of the optimal dispersal strategies depends on the ecological conditions, and we screen through a set of ecological parameters to find the relationship between ecology and optimal dispersal strategies, see section 1.5.

The model operates on several time scales. Here, we use a specific terminology to indicate, which time scale is addressed at the moment. The largest unit of simulation is a *replicate* of evolution computing an optimal dispersal strategy at a given ecological conditions. Each replicate is composed of a sequence of selection *rounds*. A selection round takes a number of competing strategies and ranks them according to the performance in the course of population dynamics. Removal of sub-optimal strategies and the introduction of novel mutants occurs between rounds. Each selection round is composed of a chain of simulation *steps*. A simulation step updates the meta-population state according to the chosen ecological conditions and operating dispersal strategies. Finally, at each step, a sequence of *events* is executed, each event representing one specific process occurring in the system, such as consumers foraging, or resource reproduction. Sections 1.1 and 1.2 cover the timescales of events and simulation steps, while the timescales of selection rounds and evolution replicates are addressed in sections 1.4 and 1.5.

#### 1.1 Informed dispersal

The very first event executed at each simulation step is the dispersal of consumers. There, each foraging consumer makes an independent decision either to stay in the current patch or leave it and move to a randomly selected patch. This decision is stochastic, i.e. two individuals with identical dispersal strategies located in the same patch may make different decisions: one will stay while another will depart. However, the rate of dispersal ($m$), i.e. the probability of leaving a patch per time unit, deterministically depends on the patch state and the strategy employed. This section presents how the dispersal rate is calculated.

We consider three sensory models different in what aspect of patch state is perceived by the consumers:

1. In the **uninformed** model, consumers do not get any information about the patch state. Here, they make an uninformed decision on the dispersal. This serves as the null model.

2. In the **conspecifics sensing** model, consumers sense the number of consumers in the current patch ($N_c$).

3. In the **resource sensing** model, consumers sense the amount of resource in the current patch ($N_r$).

All consumers sense their patch state simultaneously, and the obtained information is accurate.

In the uninformed model, the dispersal rate is independent of the patch state and is defined by the dispersal strategy of the consumer:

$$m = m_0, \tag{1}$$

where $m_0$ is the dispersal rate given by the strategy.

In the models with informed dispersal, the dispersal rate of a consumer depends on its strategy and the sensed information in a sigmoid manner

$$m(I) = L + (R - L)\left(1 + e^{-\frac{I - I_0}{\sigma}}\right)^{-1}, \tag{2}$$

where $I = N_c$ (number of consumers in the patch) or $I = N_r$ (amount of resource in the patch) depending on the sensing model, $L$ and $R$ are the limits of dispersal rate for very low and very high values of sensory input, $I_0$ is the position of the inflection point of the sigmoid, and $\sigma$ is the width of the transition between limits. Dispersal rate is capped at 0 from below, i.e. if for any reason $m(I) < 0$, a consumer abstains from dispersal (same as $m(I) = 0$).

The parameters of the dispersal strategy ($m_0$ or $L, R, I_0, \sigma$) are subjected to evolutionary adaptation, see section 1.4.

#### 1.2 Population dynamics

The population dynamics follows a stochastic, individual-based simulation of two species (resource and consumer) on a meta-population. In the course of simulation, resource species individuals reproduce, are being predated by consumers, and, rarely, immigrate from an external source (so resource-depleted patches are not completely lost and can be eventually repopulated). Consumers, in turn, spontaneously die, forage for resource, process it to reproduce, and disperse across patches. All these events occur in the fixed order during every simulation step, see Fig. S1.

**Population state description**

Each consumer in the model can be in one of two mutually exclusive states: *foraging* or *processing*. The difference between these states is that only foraging individuals can disperse among patches and forage for the resource, and only processing individuals can reproduce. Individual's state changes: a foraging individual turns into a processing one, when it acquires a resource, and a processing individual gives rise to two foraging ones upon reproduction. Any consumer individual is completely characterized by its state, location on the meta-population, and the dispersal strategy executed. For the purpose of the calculation speed, the model does not simulate each individual independently and only keeps track of the number of consumers with the same combination of these three characteristics.

Representation of the resource species is coarse-grained: each resource "individual" represents the amount of resource biomass that a single consumer needs to process in order to reproduce. A resource individual is completely characterized by its location on the meta-population. Thus, the model only keeps track on the count of resource in each patch.

At each simulation step, the model computes the change in the population state within a short time period ($\mathbf{e}$). To do so, the model implements 6 events in the following order:

**Event 1: foraging consumers disperse.**

This event begins with foraging consumers sensing the state of the patch ($I$) and evaluating their dispersal rate ($m$), see section 1.1 for details.

With the known dispersal rate, the residence time of an individual is distributed exponentially, hence the probability for a given consumer to leave the current patch during the simulation step $\mathbf{e}$ is

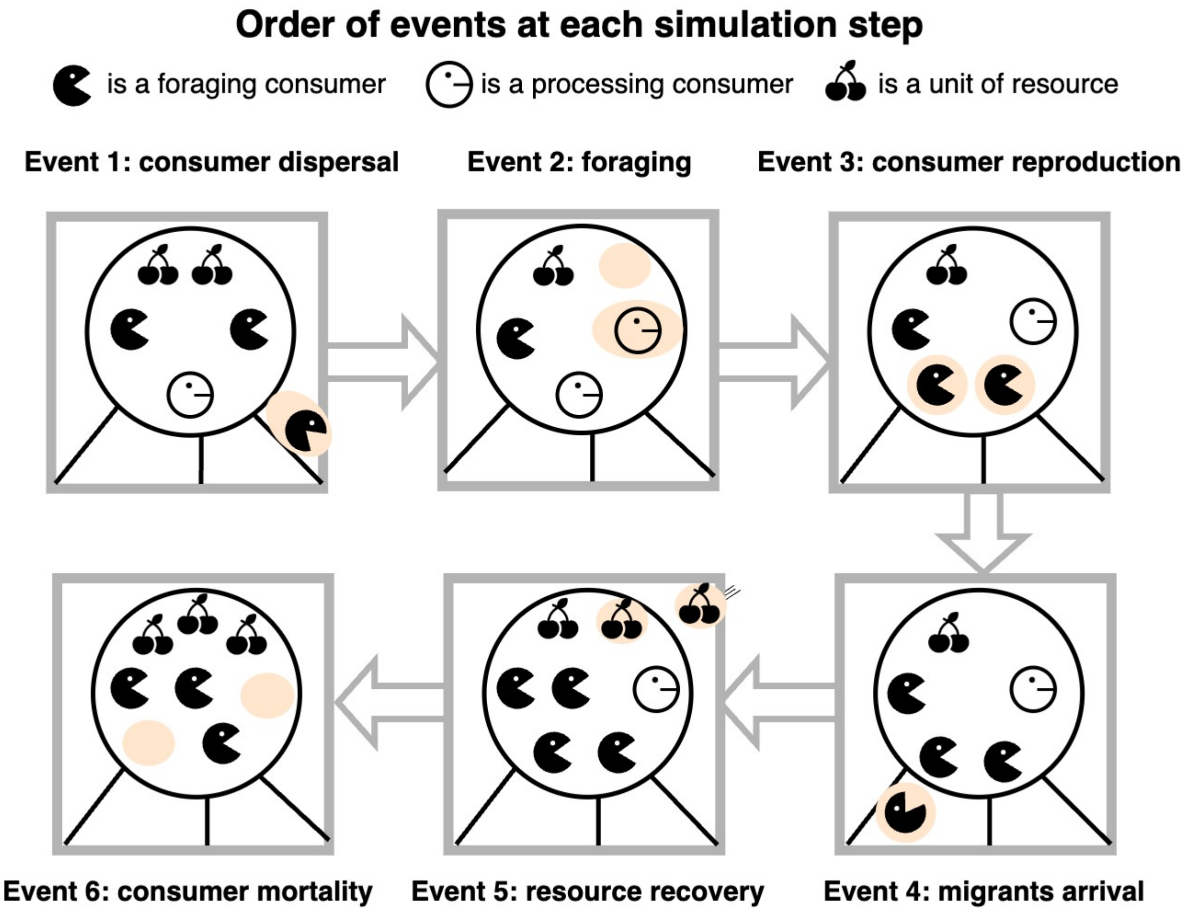

Figure S1: **Order of events in a simulation step.** Each of six frames depicts a population of a single patch (big circle) after each event. The changes from the previous frame are highlighted in color. The order shown illustrates our implementation of the model, and other conceptually equivalent orders are possible. However, the foraging (event 2) must occur between departure (event 1) and arrival (event 3) of dispersing consumers, as the dispersal would not have a costs otherwise.

equal to

$$P_{\text{dispersal}} = 1 - e^{-m\mathbf{e}}, \tag{3}$$

where $m$ is the dispersal rate and $\mathbf{e}$ is the duration of the simulation step.

Then the number of individuals dispersed during the time step is drawn as a random value with the binomial distribution, which success rate is $P_{\text{dispersal}}$ and the number of trials is equal to the number of foraging individuals in the patch who execute a given dispersal strategy - $N_{f,S}$.

$$N_{f,S,\text{dispersed}} \sim \text{Bin}(x|N_{f,S}, P_{\text{dispersal}}). \tag{4}$$

As a result of dispersal, the number of foraging individuals in the patch having a given dispersal strategy falls

$$N_{f,S} \to N_{f,S} - N_{f,S,\text{dispersed}}. \tag{5}$$

Calculations (3)-(5) are performed independently for each patch and for each dispersal strategy. The same applies to all events below.

**Event 2: foraging consumers go foraging.**

Next, the foraging consumers, which did not disperse make an attempt to secure resource. All foraging consumers forage with the rate $\omega_c$. The probability of a given resource individual to escape being captured by a given foraging consumer during one simulation step is

$$P_{\text{escape one forager}} = e^{-\omega_c \mathbf{e}}. \tag{6}$$

A resource individual survives the foraging event only if it is not captured by any of $N_f$ foragers, which happens with probability

$$P_{\text{escape all foragers}} = (P_{\text{escape one forager}})^{N_f} = e^{-\omega_c N_f \mathbf{e}}. \tag{7}$$

The number of captured resource individuals is drawn from the binomial distribution

$$N_{r,\text{consumed}} \sim \text{Bin}(x|N_r, 1 - P_{\text{escape all foragers}}), \tag{8}$$

where $N_{r,\text{consumed}}$ is the number of resource individuals acquired by foraging consumers, and $N_r$ is the total amount of resource in the patch.

As a result, $N_{r,\text{consumed}}$ of resource are lost, and the same number of randomly chosen foraging consumers change their state to the processing one.

$$\begin{aligned} N_r &\to N_r - N_{r,\text{consumed}}, \\ N_f &\to N_f - N_{r,\text{consumed}}, \\ N_p &\to N_p + N_{r,\text{consumed}}, \end{aligned} \tag{9}$$

where $N_p$ is the number of processing individuals.

Which consumers get the food and turn to processing state is chosen randomly among all foraging individuals in the patch, i.e. the difference in dispersal strategies among consumers has no effect on the foraging efficiency.

**Event 3: processing consumers reproduce.**

Following the literature tradition, the processing is characterized by the processing time ($h_c$), thus, the rate of reproduction of processing individuals is $1/h_c$. The probability of a processing consumer to reproduce within a single simulation step is

$$P_{\text{reproduction}} = 1 - e^{-\mathbf{e}/h_c}. \tag{10}$$

Thus, the number of reproduced consumers is drawn from the binomial distribution

$$N_{p,\text{reproduced}} \sim \text{Bin}(x|N_p, P_{\text{reproduction}}), \tag{11}$$

where $N_p$ is the number of processing individuals in the patch.

Then each reproduced individual turns into two foraging ones.

$$\begin{aligned} N_p &\to N_p - N_{p,\text{reproduction}}, \\ N_f &\to N_f + 2N_{p,\text{reproduction}}. \end{aligned} \tag{12}$$

The new foraging individuals inherit the dispersal strategy of the maternal individual and are put into the same patch.

**Event 4: arrival of dispersed consumers.**

Here, foraging individuals dispersed at the event 1 arrive to their destination patches. Each individual disperses stochastically and independently from others, so two identical individuals from the same patch may arrive to different destination patches. As the chosen geometry of the meta-population is the fully connected graph, an individual may arrive to any patch with equal probability.

In the result, foraging population increases by

$$N_f \to N_f + N_{f,\text{arrived}}, \tag{13}$$

where $N_{f,\text{arriving}}$ is the number of arrived individuals. For a single patch, the numbers of departing and arriving individuals are different, but each departing individual always arrives at some patch $\left(\sum_{\text{patches}} N_{f,\text{arrived}} = \sum_{\text{patches}} N_{f,\text{dispersed}}\right)$.

Dispersed consumers do not participate in foraging (event 2) and this constitutes the cost of dispersal.

**Event 5: Resource population recovers.**

This event combines three processes related to the recovery of the resource population:

1. Resource population grows logistically. The growth rate is given by $\omega_{\text{effective}} = \omega_r\,(1 - N_r/K_r)$, where $\omega_r$ is the base growth rate of the resource species, $N_r$ is the amount of resource in a patch, and $K_r$ is the resource carrying capacity.

2. Resource arrive with a small rate $E_r = 0.1$ from an external source. The value is small enough, so that consumer population cannot be sustained by this inflow alone (see also Section 1.3 below).

3. The amount of resource is capped at the carrying capacity $K_r$.

The number of resource individuals emerged due to growth and arrival are random numbers sampled from Poisson distributions:

$$N_{r,\text{growth}} \sim \text{Poisson}(x|\omega_{\text{effective}} N_r \mathbf{e}), \tag{14}$$

$$N_{r,\text{immigration}} \sim \text{Poisson}(x|E_r \mathbf{e}). \tag{15}$$

As the result, the number of prey in a patch changes to

$$N_r \to \min\left(N_r + N_{r,\text{growth}} + N_{r,\text{immigration}}, K_r\right). \tag{16}$$

**Event 6: Consumers fall to intrinsic mortality.**

All consumers are subjected to the intrinsic mortality with the same rate $d_c$. The probability of a given consumer to die at a single step is given by

$$P_{\text{death}} = 1 - e^{-d_c \mathbf{e}}, \tag{17}$$

This way, the number of individuals dying at the step is a random value drawn from the binomial distribution

$$N_{f,\text{death}} \sim \text{Bin}(x|N_f, P_{\text{death}}), \tag{18}$$

$$N_{p,\text{death}} \sim \text{Bin}(x|N_p, P_{\text{death}}), \tag{19}$$

Then the numbers of foraging and processing individuals reduce:

$$N_f \to N_f - N_{f,\text{death}}, \tag{20}$$

$$N_p \to N_p - N_{p,\text{death}}. \tag{21}$$

This is the last event in a simulation step. Once this event concludes, the next simulation step immediately begins, where the same events occur again in the same order.

#### 1.3 Stability analysis of the population dynamics in an isolated patch

In the limit of large resource and consumers population, we can consider the analytical model of population dynamics in an isolated patch. Such a model is helpful to describe the "base line" of the population dynamics in the absence of dispersal.

If we denote the number of resource individuals, foraging and processing consumers in the patch as $N_r$, $N_f$, and $N_p$, respectively, the dynamics of a patch state is governed by a set of ordinary differential equations

$$\begin{aligned} \frac{d}{dt}N_r &= \omega_r N_r \left(1 - \frac{N_r}{K_r}\right) - \omega_c N_f N_r, \\ \frac{d}{dt}N_f &= -\omega_c N_f N_r - d_c N_f + \frac{1 + r_c}{h_c} N_p, \\ \frac{d}{dt}N_p &= \omega_c N_f N_r - d_c N_p - \frac{1}{h_c} N_p, \end{aligned} \tag{22}$$

where $\omega_r$ is the growth rate of resource, $K_r$ is the carrying capacity of resource, $\omega_c$ is the foraging rate of consumers, $d_c$ is the intrinsic mortality rate of consumers, and $h_c$ is the handling time of consumers, and $r_c$ is the number of foraging individuals produced in the act of reproduction (in addition to the maternal individual).

In our numerical model, the number of offspring produced by a processing individual is always $r_c = 1$ but in this analytical section, we allow this parameter to vary.

This system has three stationary points:

1. Stationary point with an empty population,

$$N_f^* = 0, \quad N_p^* = 0, \quad N_r^* = 0. \tag{23}$$

2. Stationary point with resource-only population,

$$N_f^* = 0, \quad N_p^* = 0, \quad N_r^* = K_r. \tag{24}$$

3. Stationary point with coexisting resource and consumers,

$$\begin{aligned} N_f^* &= \frac{\omega_r}{\omega_c}(1-F), \\ N_p^* &= \frac{\omega_r}{\omega_c}(1-F)\frac{Yr_c - 1}{1+r_c}, \\ N_r^* &= FK_r, \end{aligned} \tag{25}$$

where

$$Y = \frac{1 + h_c d_c}{r_c - h_c d_c}. \tag{26}$$

is the consumer processing factor and

$$F = \frac{Y d_c}{\omega_c K_r} = \frac{d_c}{\omega_c K_r}\frac{1 + h_c d_c}{r_c - h_c d_c} \tag{27}$$

is the fraction of resource carrying capacity surviving at equilibrium. Note that the values of $Y$ are either $Y > 1/r_c$ if $h_c d_c < r_c$, or $Y < -1$ otherwise.

**Existence and stability of stationary points**

Stationary point 1 (empty population) exists at any combination of positive values of ecological parameters $(\omega_c, h_c, r_c, d_c, \omega_r, K_r)$ but is always unstable.

Stationary point 2 (resource-only population) also always exists and is unstable only if two conditions are satisfied: $F < 1$ and $Y > 0$. Or, in terms of the raw ecological parameters

$$\begin{aligned} &r_c > h_c d_c, \text{ and} \\ &\omega_c K_r \frac{r_c - h_c d_c}{1 + h_c d_c} > d_c. \end{aligned} \tag{28}$$

The first inequality here means that the mortality during the processing stage is lower than the reproduction rate achieved there. In other words, entering the processing mode causes a positive population change. The second inequality states that in the best conditions (resource is at the carrying capacity), consumers reproduce faster than they die out.

Stationary point 3 (mixed resource/consumer population) is biologically meaningful $(N_f^*, N_p^*, N_r^* > 0)$ only if $F < 1$ and $Y > 1/r_c$. Hence, the stability of resource-only state (24) is mutually exclusive with the existence of the mixed population (25).

The equilibrium with mixed population, given by the stationary point 3, may also be unstable. Stability is achieved when all real parts of the eigenvalues of the Jacobian of Eq. (22) are negative.

The eigenvalues are given by the roots of the cubic characteristic equation

$$\begin{aligned}
&c_3\lambda^3 + c_2\lambda^2 + c_1\lambda + c_0 = 0, \text{ where}\\
&\quad c_3 = Y(Yr_c - 1),\\
&\quad c_2 = FK_r\omega_c(Y^2 r_c + 2Yr_c - 1) + FY\omega_r(Yr_c - 1),\\
&\quad c_1 = FK_r\omega_r\omega_c(Y - F(Y+1) + Yr_c(2F(Y+1) - Y)),\\
&\quad c_0 = F^2 K_r^2 \omega_c^2 \omega_r (1 + r_c)(1 - F).
\end{aligned} \tag{29}$$

Routh-Hurwitz criterion states that the real parts of all roots are negative if $c_2/c_3 > 0$, $c_1/c_3 > 0$, $c_0/c_3 > 0$, and $c_2c_1/c_3^2 > c_0/c_3$. Since the existence of the meaningful stationary point requires $F < 1$ and $Y > 1/r_c$, this implies $c_3, c_2, c_0 > 0$. This makes two of the four Routh-Hurwitz inequalities to be automatically satisfied. Between the remaining two inequalities, the inequality $c_2c_1/c_3^2 > c_0/c_3$ is strictly stronger than $c_1/c_3 > 0$ and leads to the following stability condition:

$$\begin{aligned}
&a_4Y^4 + a_3Y^3 + a_2Y^2 + a_1Y + a_0 > 0, \text{ where}\\
&\quad a_4 = (2F - 1)r_c^2(K_r\omega_c + \omega_r),\\
&\quad a_3 = ((6r_c - 1)F - 2r_c + 1)K_r\omega_c - ((3 - 2r_c)F - 2)r_c\omega_r,\\
&\quad a_2 = ((5r_c - 4)F + 2 - r_c)r_cK_r\omega_c - ((3r_c - 1)F + 1)\omega_r,\\
&\quad a_1 = -(5F - 1)r_cK_r\omega_c + F\omega_r,\\
&\quad a_0 = FK_r\omega_c.
\end{aligned} \tag{30}$$

If the condition (30) is satisfied, the stationary point (25) is an attracting stable point, otherwise, the population executes a limit cycle around it.

**Correlation between the survived fraction of resource and the stability of the equilibrium**

The stability of equilibrium is determined by the sign of the largest real part of the Jacobian eigenvalues (29). However, we found that the fraction of resource carrying capacity at equilibrium ($F = N_r^*/K_r$) is highly correlated with the stability of the equilibrium population, see Fig. 2C in the main text. The decision tree classifier trained later on the viability data shows that the fraction of survived resource $F$ a better feature for distinguishing viable ecologies from non-viable ones than the largest real part of eigenvalues.

**Resource immigration ($E_r$) impact**

Next, we consider how the equilibrium is affected by the immigration of resource – the process used to repopulate patches with extinct resource population. If there is an inflow of resource individuals with rate $E_r$, the population dynamics equation system (22) becomes

$$\begin{aligned}
\frac{d}{dt}N_r &= \omega_r N_r \left(1 - \frac{N_r}{K_r}\right) - \omega_c N_f N_r + E_r,\\
\frac{d}{dt}N_f &= -\omega_c N_f N_r - d_c N_f + \frac{2}{h_c}N_p,\\
\frac{d}{dt}N_p &= \omega_c N_f N_r - d_c N_p - \frac{1}{h_c}N_p.
\end{aligned} \tag{31}$$

This system has three stationary points:

1. Unreachable stationary point with negative resource population,

$$N_f^* = 0, \quad N_p^* = 0, \quad N_r^* = -\frac{K_r}{2}\left(\sqrt{1+\frac{4E_r}{\omega_r K_r}} - 1\right). \tag{32}$$

$$\tag{33}$$

2. Stationary point with resource-only population,

$$N_f^* = 0, \quad N_p^* = 0, \quad N_r^* = \frac{K_r}{2}\left(\sqrt{1+\frac{4E_r}{\omega_r K_r}} + 1\right). \tag{34}$$

$$\tag{35}$$

3. Stationary point with coexisting resource and consumers,

$$\begin{aligned} N_f^* &= \frac{\omega_r}{\omega_c}(1-F) + \frac{E_r}{Y d_c}, \\ N_p^* &= \left(\frac{\omega_r}{\omega_c}(1-F) + \frac{E_r}{Y d_c}\right)\frac{Y r_c - 1}{1+r_c}, \\ N_r^* &= F K_r, \end{aligned} \tag{36}$$

In the mixed population solution, the number of resource individuals at equilibrium is independent on the strength of the immigration flow $E_r$. However, the equilibrium consumer population grows with an increase in inflow. This opens an opportunity for a consumer population to survive on the resource immigration inflow alone. Such a dynamics would contradict the focus of this study, and we are interested in the conditions, where this does not happen.

Eqs. (36) show that the impact of the resource immigration on the equilibrium number of consumers is additive. The total equilibrium consumer population ($N_f^* + N_p^*$) is increased by

$$\Delta_{N_f^* + N_p^*} = \frac{E_r r_c}{d_c}\frac{1}{1+r_c}\left(1+\frac{1}{Y}\right) < \frac{E_r r_c}{d_c}, \tag{37}$$

where the last inequality immediately follows from the condition $Y > 1/r_c$, which is necessary for an existence of mixed population, see Fig. (25). Hence, the consumer population increase is of the order of magnitude of $\frac{E_r r_c}{d_c}$. To make sure that the immigration of resource is insufficient to sustain the consumers population, it is sufficient to satisfy the condition $\frac{E_r r_c}{d_c} \ll 1$. In our work we use $r_c = 1$, $d_c = 1$ and $E_r = 0.1$, such that $\frac{E_r r_c}{d_c} = 0.1$ always.

#### 1.4 Evolutionary dynamics

Evolution of dispersal strategies is implemented as an iteration of mutation and selection processes, in which new strategies are introduced and then the strategies with lower performance are removed from population.

**Initialization of the set of competing strategies**

In each replicate of evolution, the initial population is an independently drawn sample of 10 random dispersal strategies (see section 1.1 for strategy definition).

In the uninformed model, the initial dispersal rates values $m_0$ were drawn from log-uniform distribution:

$$\log_{10}(m_0) \sim U(-2, 2), \tag{38}$$

where $U(a, b)$ means a uniform distribution with values drawn from the interval $(a, b)$.

In the resource-sensing model, all four parameters were drawn from log-uniform distributions:

$$\begin{aligned} \log_{10}(L) &\sim U(0, 2), \\ \log_{10}(R) &\sim U(-2, 0), \\ \log_{10}(I_0/N_r^*) &\sim U(-2, 0), \\ \log_{10}(\sigma/N_r^*) &\sim U(-2, 0), \end{aligned} \tag{39}$$

where $N_r^*$ is the equilibrium amount of resource according to the analytical model (given by Eq. (25) in section 1.3).

In the conspecifics sensing model, the initial sampling is almost the same. The only differences is that the $L$ and $R$ sampling distributions are swapped (because conditions promoting dispersal are resource amount being low but consumer numbers being high) and that the $I_0$ and $\sigma$ are scaled to the equilibrium number of consumers $N_c^*$:

$$\begin{aligned} \log_{10}(L) &\sim U(-2, 0), \\ \log_{10}(R) &\sim U(0, 2), \\ \log_{10}(I_0/N_c^*) &\sim U(-2, 0), \\ \log_{10}(\sigma/N_c^*) &\sim U(-2, 0). \end{aligned} \tag{40}$$

**Selection round simulation**

Each selection round begins with $A = 10$ competing dispersal strategies. The meta-population is initialized with each patch having $N_r^*$ resource individuals in it (rounded up), and a number of consumers randomly distributed among all patches. The total number of consumers in either foraging or processing states is initialized to be equal to the number there would be at the analytical equilibrium (Eq. (25)) across all patches. Each competing strategy has an equal share in both foraging and processing pools of initial consumers, so if there are $A$ strategies in a meta-population of $M$ patches, in the ecology with $N_f^*$ foraging consumers at equilibrium, each strategy starts with $MN_f^*/A$ foraging individuals (rounded up). The initial number of processing individuals follows the same rule. However, this pool of individuals is randomly assigned to patches, so the patches themselves are not necessarily at the equilibrium.

Then, this population follows the population dynamics described in section 1.2. During this dynamics, some dispersal strategies may go extinct. Selection round lasts either for 5000 simulation steps, or until only 5 strategies remain, whichever condition is satisfied first. After the end of the selection round, the 5 most numerous (in terms of their cumulative population size $N_c + N_f$) dispersal strategies are sampled to continue evolution, and remaining strategies are discarded.

**Mutation protocol**

Each selection round ends with 5 strategies being selected. Then, before the next selection round starts, each of these strategies produces a single mutant. Parameters of the mutant decision function are based on the parental strategy but are randomly disturbed.

In the uninformed model, the mutant dispersal rate ($m'_0$) is formed as

$$\log_{10}(m'_0) = \log_{10}(m_0) + \delta m_0, \quad \delta m_0 \sim N(x|0, 0.3), \tag{41}$$

where $\delta m_0$ is a random value sampled from a normal distribution with mean zero and variance 0.3 (i.e. the value changes by about 30%).

In the resource sensing and consumer sensing models, the parameters of mutant are computed similarly

$$\begin{aligned} \log_{10}(L') &= \log_{10}(L) + \delta L, \quad \delta L \sim N(x|0, 0.3), \\ \log_{10}(R') &= \log_{10}(R) + \delta R, \quad \delta R \sim N(x|0, 0.3), \\ \log_{10}(I'_0) &= \log_{10}(I_0) + \delta I_0, \quad \delta I_0 \sim N(x|0, 0.3), \\ \log_{10}(\sigma') &= \log_{10}(\sigma) + \delta\sigma, \quad \delta\sigma \sim N(x|0, 0.3). \end{aligned} \tag{42}$$

To prevent the strategies from getting stuck at the extreme values, where selection is insensitive to mutations, the values of produced mutants were limited by the intervals:

$$\begin{aligned} L' &\in [10^{-3}, 10^3], \\ R' &\in [10^{-3}, 10^3], \\ I_0 &\in [\min(1, I^*/5), 10I^*], \\ \sigma &\in [\min(1, I^*/5), 10I^*], \end{aligned} \tag{43}$$

where $I^* = N_r^*$ or $I^* = N_c^*$ - the equilibrium value of the sensed information.

The 5 winners of the previous selection rounds and the 5 mutants comprise the set of 10 dispersal strategies competing in the next selection round.

**Identifying the optimal dispersal strategy**

A complete evolutionary replicate consists of 100 selection rounds. At the end of the last round, the single most numerous strategy is picked as the endpoint of evolution.

For each combination of ecological parameters, we simulate 10 independent replicates of evolution. From each replicate we take a single endpoint strategy. After that, these 10 endpoint strategies compete among each other in the final selection round to determine the overall best strategy. This final round is independently repeated 100 times (with the same set of competing strategies) and each time, a single most numerous strategy is identified. The strategy with the highest win rate in this series is recorded as an optimal dispersal strategy for the given ecological parameters.

#### 1.5 Ecology screening

We found optimal dispersal strategies for a wide range of different ecologies. An ecology is determined by six parameters: $\omega_c, h_c, d_c, \omega_r, K_r, E_r$. Without loss of generality, we set the time scale of the population dynamics by using $d_c = 1$. To ensure that the inflow of resource is small enough to not influence the population dynamics of consumers, we set $E_r = 0.1$, see the end of the section 1.3 for details. This leaves 4-dimensional parameter space of ecologies. We randomly sampled the space, with parameter values drawn as following:

$$\begin{aligned} \log_{10}(\omega_c) &\sim U(-3, -1), \\ \log_{10}(\omega_r) &\sim U(-2, 1), \\ \log_{10}(K_r) &\sim U(2, 3.5), \\ h_c &\sim U(0, 1). \end{aligned} \tag{44}$$

Preliminary simulations have shown that ecologies with $N_c^* < 3$ do not evolve strategies capable to support a viable population. To improve the efficiency of our simulations, we did not simulate evolution of these ecologies but still recorded them as non-viable conditions.

### 2 Knockout competition

We tested the evolved strategies against a number of knockout strategies. Tests against each knockout were performed independently. In each test, we run a simulation round of population dynamics with a meta-population, which initially contained both the evolved strategy and one of knockouts in equal proportions. Each evolved strategy has been tested in the same ecology, in which it evolved at the first place. The strategy with more individuals at the end of the simulation round was declared a winner. The simulation round was repeated 100 times, and the win rate of the evolved strategy against a given knockout was recorded.

Knockouts were implemented as sigmoid decision functions with modified parameters. For each tested strategy, the following knockouts are used:

**Knockout 1: no dispersal**

The first knockout does not disperse at all. If the parameters of the evolved decision function are $L, R, \sigma, I_0$, then the parameters of the first knockout are:

$$\begin{aligned} L_1 &= 0, \\ R_1 &= 0, \\ \sigma_1 &= \sigma, \\ I_{0,1} &= I_0. \end{aligned} \tag{45}$$

Note that since the knockout function is constant, the values of $\sigma_1$ and $I_{0,1}$ have no impact, see Fig. S2.

**Knockout 2: low limit insensitive**

The low limit insensitive knockout always disperses at the minimally experienced dispersal rate $\min(m(I))$, obtained from the the population dynamics of evolved strategy alone, and is formally defined as a sigmoid function with parameters equal to

$$\begin{aligned} L_2 &= \min(m(I)), \\ R_2 &= \min(m(I)), \\ \sigma_2 &= \sigma, \\ I_{0,2} &= I_0. \end{aligned} \tag{46}$$

**Knockout 3: high limit insensitive**

This knockout is similar to the low limit insensitive but uses the highest recorded dispersal rate instead – $\max(m(I))$:

$$\begin{aligned} L_3 &= \max(m(I)), \\ R_3 &= \max(m(I)), \\ \sigma_3 &= \sigma, \\ I_{0,3} &= I_0. \end{aligned} \tag{47}$$

see Fig. S2

**Knockout 4: no baseline dispersal**

This knockout tests the dispersal dynamics in a favorable patch.

To do so, this knockout strategy keeps the position of the inflection point $I_0$, the transition scale $\sigma$, and the maximal experienced dispersal rate $\max(m(I))$ of the original evolved strategy, but drops the minimal experienced dispersal rate to zero: $\min(m_4(I)) = 0$, Fig. S2.

In the conspecifics sensing model, where an underpopulated patch is a favorable condition,

$$\begin{aligned}
L_4 &= \frac{m(\max(N_c))}{1 - \frac{F_L}{F_R}}, \\
R_4 &= L_4(1 - F_L), \\
\sigma_4 &= \sigma, \\
I_{0,4} &= I_0, \\
&\text{where,} \\
F_L &= 1 + e^{-(\min(N_c) - I_0)/\sigma}, \\
F_R &= 1 + e^{-(\max(N_c) - I_0)/\sigma},
\end{aligned} \tag{48}$$

while $\min(N_c)$ and $\max(N_c)$ are the minimal and maximal number of consumers experienced in the preliminary simulation round (with evolved strategy alone).

In the resource sensing model, where a resource-rich patch is a favorable condition,

$$\begin{aligned}
L_4 &= \frac{m(\min(N_r))}{1 - \frac{F_R}{F_L}}, \\
R_4 &= L_4(1 - F_R), \\
\sigma_4 &= \sigma, \\
I_{0,4} &= I_0, \\
&\text{where,} \\
F_L &= 1 + e^{-(\min(N_r) - I_0)/\sigma}, \\
F_R &= 1 + e^{-(\max(N_r) - I_0)/\sigma},
\end{aligned} \tag{49}$$

while $\min(N_r)$ and $\max(N_r)$ are the minimal and maximal amount of resource experienced in the preliminary simulation round (with evolved strategy alone).

**Knockout 5: no saturation**

This knockout tests the tolerance of the dispersal strategy to the unfavorable state of the current patch.

This knockout effectively replaces the saturating part of the sigmoid function (where $m''(I) < 0$) with the exponential growth of the dispersal rate continuing the accelerating pattern of the tested sigmoid (where $m''(I) > 0$), see Fig. S2. The resulting exponent coincide with the tested sigmoid at two critical points: at the median of the sensed information (median$(I)$) and at the point, where the population experienced the largest inclination of the decision function $I_{\text{max inc.}}$. Typically, $I_{\text{max inc.}} = I_0$, however, if $I_0$ is outside of the experienced interval of the sensed information $(\min(I), \max(I))$, then $I_{\text{max inc.}}$ is the nearest limit of that interval:

$$I_{\text{max inc.}} = \min[\max[I_0, \min(I)], \max(I)] = \text{clip}[I_0, (\min(I), \max(I))] \tag{50}$$

If we denote the dispersal rates (see Eq. (2)) at these reference points as

$$\begin{aligned} m_{\text{median}} &= L + (R - L)\left(1 + e^{-\frac{\text{median}(I) - I_0}{\sigma}}\right)^{-1}, \\ m_{\text{max inc.}} &= L + (R - L)\left(1 + e^{-\frac{I_{\text{max inc.}} - I_0}{\sigma}}\right)^{-1}, \end{aligned} \tag{51}$$

then the parameters of the unbounded knockout for the conspecifics sensing model were computed as

$$\begin{aligned} L_5 &= 0, \\ R_5 &= 10^{10} \\ \sigma_5 &= \frac{I_{\text{max inc.}} - \text{median}(I)}{\ln\left(\frac{m_{\text{max inc.}}}{m_{\text{median}}}\right)}, \\ I_{0,5} &= I_{\text{max inc.}} - \sigma_5 \ln\left(\frac{m_{\text{max inc.}}}{R_5}\right), \end{aligned} \tag{52}$$

and for the resource sensing model as

$$\begin{aligned} L_5 &= 10^{10}, \\ R_5 &= 0 \\ \sigma_5 &= \frac{I_{\text{max inc.}} - \text{median}(I)}{\ln\left(\frac{m_{\text{median}}}{m_{\text{max inc.}}}\right)}, \\ I_{0,5} &= I_{\text{max inc.}} - \sigma_5 \ln\left(\frac{L_5}{m_{\text{max inc.}}}\right). \end{aligned} \tag{53}$$

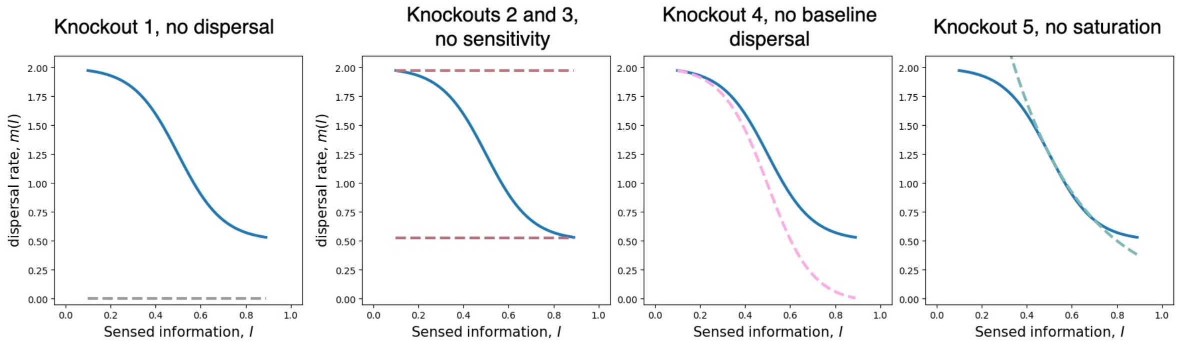

Figure S2: **Knockout modification of an example decision function.** Each panel shows a sample decision function (blue) together with a knockout(s) computed according to Eqs. (45) - (53). Parameters of the sample decision function are: $L = 2$, $R = 0.5$, $I_0 = 0.5$, $\sigma = 0.1$, and for illustrative purposes, we assume the population dynamics features $\min(I) = 0.1$, $\text{median}(I) = 0.7$, $\max(I) = 0.9$.

### 3 Viability of consumer populations across ecologies and sensory models

The selection protocol employed in our study (see section 1.4) finds the most successful dispersal strategy but does not assess whether it can support a viable consumers population. For instance, if at any moment of evolution, all competing strategies drive the population towards extinction, the selection round will pick those five strategies, which survive longer than others and pass them all to the next round. The selection process favors dispersal strategies, which keep the population alive for a longer stretch of time, but there is no guarantee that the strategy obtained at the end of the evolutionary simulations is able to avoid consumers extinction.

To measure the viability of a strategy evolved in a given ecology, we simulate the population dynamics for the duration of the single selection round (5000 simulation steps). For each ecology, we repeat this simulations 100 times, and found the fraction of simulation runs, at which the consumer population stayed alive until the end of the round. The distribution of viabilities demonstrates that in most of the ecologies, population is either always go extinct or always survive until the end of the selection round see Fig. S3. Only in a few ecologies (less then 15% of all ecologies in each model), the evolved strategies produce a population of consumers, which may survive until the end of the round but is not guaranteed to do so, see Table 1. We define a *viable* ecology as an ecology, at which the meta-population executing the evolved dispersal strategy survives for the duration of the selection round with probability exceeding 90%. Otherwise, we call it *non-viable* ecology.

Table 1: Number of viable and non-viable ecologies in different sensory models.

| **Sensory model** | **Always go extinct** | **Sometimes survive** | **Always survive** | **Viable** | **Non-viable** |
|---|---|---|---|---|---|
| Uninformed | 9309 (68%) | 1768 (13%) | 2524 (19%) | 2846 (21%) | 10755 (79%) |
| Conspecifics | 9134 (67%) | 1705 (13%) | 2762 (20%) | 3095 (23%) | 10506 (77%) |
| Resource | 8521 (63%) | 1470 (11%) | 3610 (26%) | 3925 (29%) | 9676 (71%) |

We find that the viable and non-viable strategies can be separated well on 2-dimensional projection of the 4-dimensional ecology parameter space. The two most important features being the number of consumers at equilibrium ($N_c^*$) and the fraction of resource survived at equilibrium ($N_r^*/K_r$). We found the most plausible borders between viable and non-viable ecologies with these features by optimizing the F-score – the harmonic mean of the precision (what fraction of ecologies predicted to be viable are actually viable) and the recall (what fraction of viable ecologies is predicted to be viable) of the classifier, see Table 2 and Fig. 2 D-E in the main text.

Table 2: The thresholds of viability in different sensory models and the accuracy of the classification.

| **Model** | Border $N_c^*$ | Border $N_r^*/K_r$ | Precision | Recall | F-score |
|---|---|---|---|---|---|
| Isolated patch | 62 | 0.26 | 0.70 | 0.85 | 0.76 |
| Uninformed | 9.9 | 0.13 | 0.80 | 0.86 | 0.83 |
| Conspecifics sensing | 8.5 | 0.11 | 0.79 | 0.86 | 0.83 |
| Resource sensing | 10.0 | 0.0 | 0.78 | 0.92 | 0.85 |

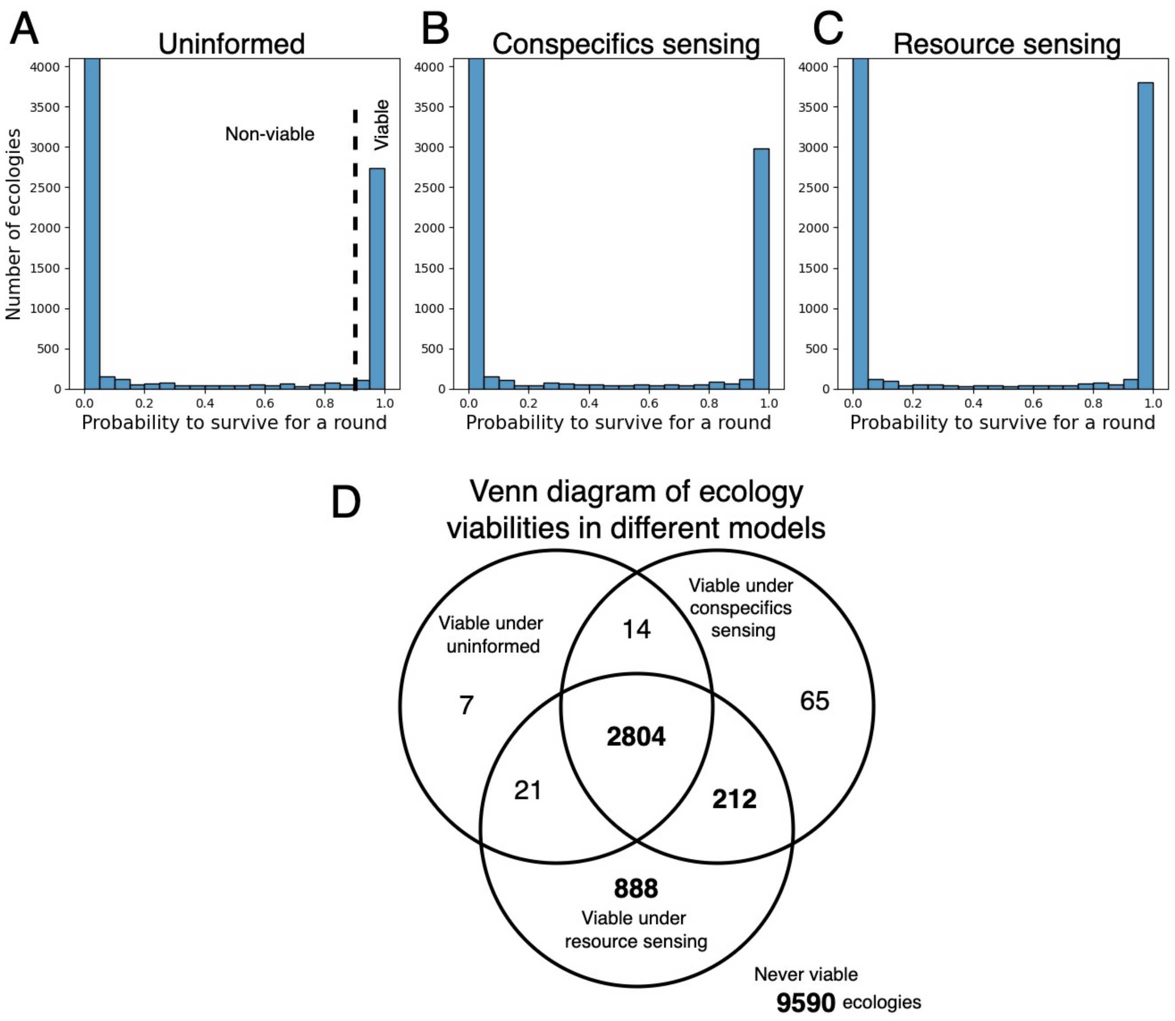

Figure S3: **Evolved dispersal strategies mostly result in perfectly viable or completely non-viable populations.** Each panel shows a histogram of survival probabilities for optimal dispersal strategies in all ecologies within a single sensory model: uninformed (**A**), conspecifics sensing (**B**), or resource sensing (**C**). We define a viable ecology / dispersal strategy, if the survival rate exceeds 90% (dashed line). **D** Venn diagram shows how many ecologies are viable in each model and their combinations.

§

### 4 The same kinds of strategies evolve in metapopulations of different sizes

Our main body of investigation studies the evolution of dispersal in the meta-population of the specific size $M = 20$ patches. In this section, we present the outcomes of evolution for meta-population of different sizes, specifically for $M = 2$ see Fig. S4, and $M = 50$ see Fig. S5.

In the calculation of the $M = 50$ map, we computed only a single replicate for each ecology, instead of 10 in our main simulation. If there are multiple local optima of the dispersal strategies, they are more likely to appear in the map on Fig. S5 compared to our main investigation. On the other hand, for this calculation, we relaxed the threshold of ecologies automatically deemed non-viable from $N_c^* = 3$ to $N_c^* = 1$, as the larger meta-population makes the survival possible for smaller patch populations.

Comparing the results in the minimal meta-population with $M = 2$ with our main results ($M = 20$), we find that the viability of the minimal meta-population is much more constrained than the larger one. The regions of viable ecologies in both sensory models are close to one found in the isolated patch, see Fig. S4E,F here and Fig.3D in the main text. Naturally, most of the viable ecologies promote no-dispersal strategies, which is the dominant class there. Nevertheless, all other classes still appear and in the same regions as in our main results. However, their former regions of optimality are often non-viable, so the presence of alternative classes is smaller.

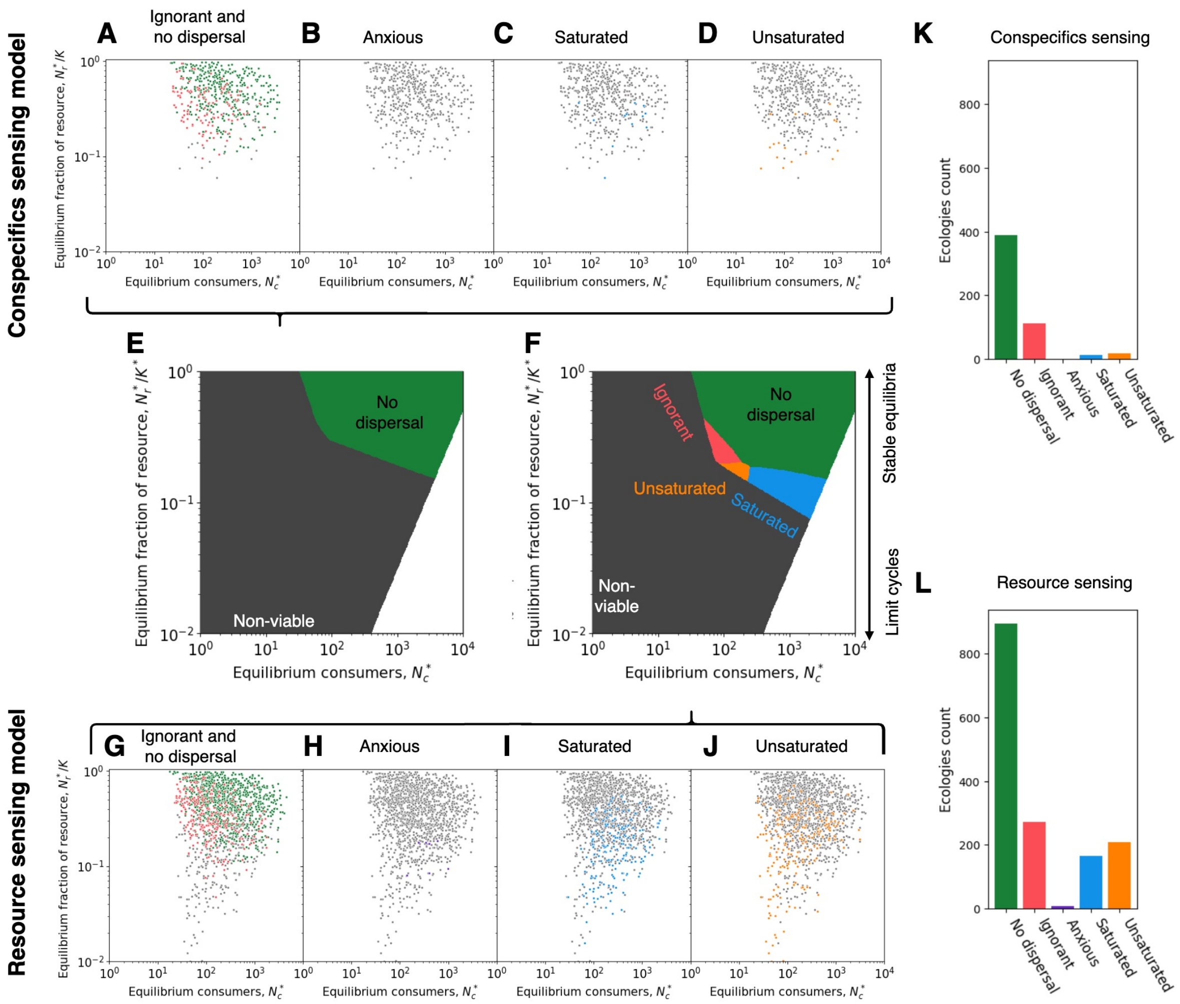

Figure S4: **Distribution of different classes of dispersal strategies in ecology space in the minimal meta-population with $M = 2$ patches. A-D** show classes evolved in the conspecifics sensing model. Each dot is a viable ecology, and the ecologies with a given class are highlighted with color. **G-J** show the same in the resource sensing model. **E, F** show the regions of optimality interpolated from these data with neural network classifier. Parameters used: $M = 2$, $d_c = 1.0$, $E_r = 0.1$, while $\omega_c, h_c, \omega_r, K_r$ vary among samples and $m_0, L, R, \sigma, I_0$ use evolved values.

In a larger meta-population with $M = 50$ patches, the regions occupied by different classes are very close to the ones observed in our main results ($M = 20$), cf. Fig. S5E,F here with Fig.5E,F in the main text. The main difference between the two maps is that the ignorant strategies are found to be optimal in ecologies with stable equilibria and low equilibrium consumers. These are the ecologies with about a dozen consumers per patch at equilibrium, such that extinctions of patches due to stochastic fluctuations are frequent and hence the leaving a patch can be always beneficial as the migrants may recolonize a patch with no competitors at all. In our main set of results, this region of the ecology space is non-viable (the critical population to survive is about 10 consumers per patch at equilibrium, see Table 1), but we still find occasional ignorant strategies there, see Fig 5A,G in the main text.

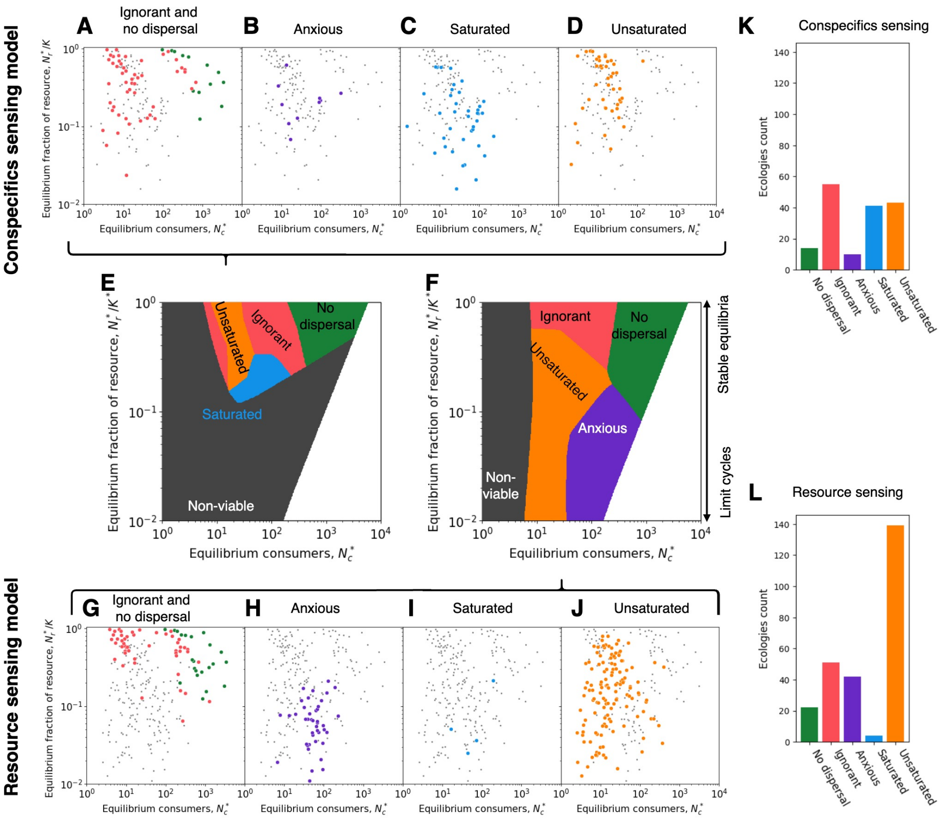

Figure S5: **Distribution of different classes of dispersal strategies in ecology space in the meta-population with $M = 50$ patches. A-D** show classes evolved in the conspecifics sensing model. Each dot is a viable ecology, and the ecologies with a given class are highlighted with color. **G-J** show the same in the resource sensing model. **E, F** show the regions of optimality interpolated from these data with neural network classifier. Parameters used: $M = 50$, $d_c = 1.0$, $E_r = 0.1$, while $\omega_c, h_c, \omega_r, K_r$ vary among samples and $m_0, L, R, \sigma, I_0$ use evolved values.

### 5 Evolution of dispersal rate in the uninformed model

In the uninformed model, the dispersal strategy is determined by a single parameter $m_0$ - the constant dispersal rate. This rate is still subjected to the evolution, and the results are shown on Fig. S6 and S7. Ecologies with larger and more stable equilibria promote lower dispersal rate. Here, our results confirm the findings that the high dispersal rate evolves under higher risk of extinctions, either due to a low population size or by high oscillations of the population dynamics.

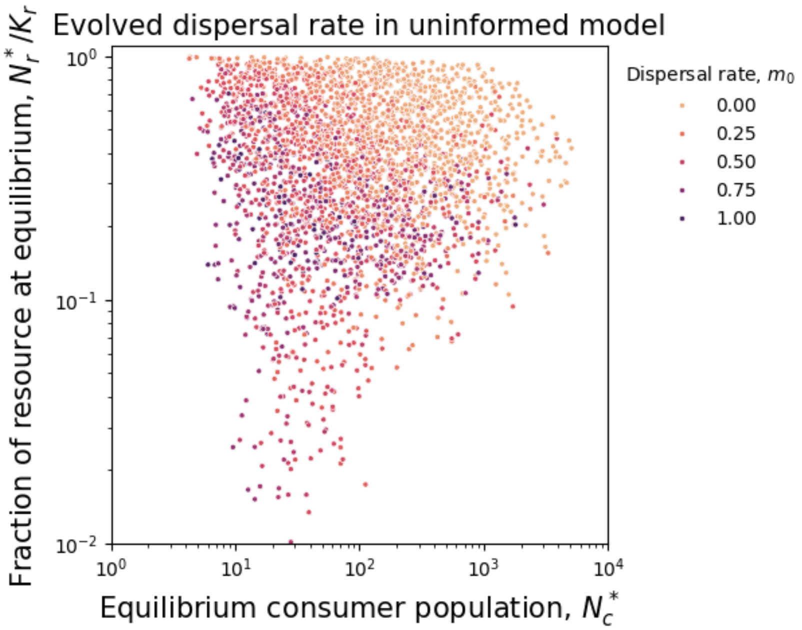

Figure S6: **Uninformed decisions promote high dispersal rate under the risk of extinction.** Each dot is a sampled ecology, with its coordinates given by its ecological parameters. The color of each dot shows the dispersal rate evolved in this sample in uninformed model. Low dispersal rates are observed in the stable regions with larger populations - where extinction is unlikely. High dispersal rates are found in the regions with low equilibrium population size and with high amplitudes of oscillations - where extinction is more likely. Non-viable populations are not shown.

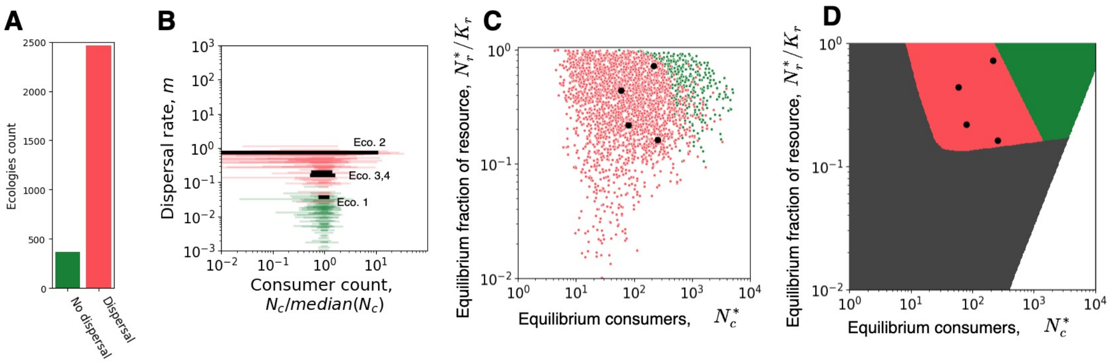

Figure S7: **Classification in uninformed model, limited to *No dispersal* and *Ignorant* classes.** Panel **A**: Histogram of the number of viable ecologies promoting each class. Panel **B**: Collection of decision functions, colored by class. Black lines show strategies evolved in the representative ecologies. Panel **C**: Distribution of classes in 2D projection of the ecological space. Each dot is an ecology, its color represents the class of evolved strategy. Panel **D**: Territories of the classes in the 2D projection of the ecological space. Red corresponds to dominance of *Ignorant* class, green - *No-dispersal*, black - non-viable ecologies, white - the region is not sampled.

### 6 Classification of the decision functions

We classify the strategies by their performance in a competition against knockouts. A meta-population is initialized with two competing strategies, the evolved one and the one of "knockouts", at equal proportions. The population dynamics is simulated for the duration of a single selection round and the strategy having more individuals at the end is recorded. This competition was independently replicated 100 times. The win rate of the evolved strategy is calculated as the fraction of replicates where it won. If the win rate of an evolved strategy is close to 100%, then the knocked out feature is adaptively significant. If the win rate is close to 50%, then the feature is not significant for the performance of the strategy. We put the threshold of the feature significance halfway between the two: at 75%.

We define the **No-dispersal** class of strategies as these, which have the win rate below 75% against the knockout number 1 – the one without dispersal, see Section 2 for knockouts definition, see Fig. S8. Among others, we define the **Ignorant** class of strategies as these, which have the win rate below 75% against either of the non-sensitive knockouts (number 2 and 3). At least one low win rate is enough. From the residual strategies, the **Anxious** strategies are defined as these, which have the win rate against knockout without base line dispersal (number 4) above 75%. The remaining strategies are separated by their win rate against the knockout without saturation (number 5): if the win rate exceeds 75%, then the strategy belongs to the **Saturated** class, and otherwise, to the **Unsaturated** class.

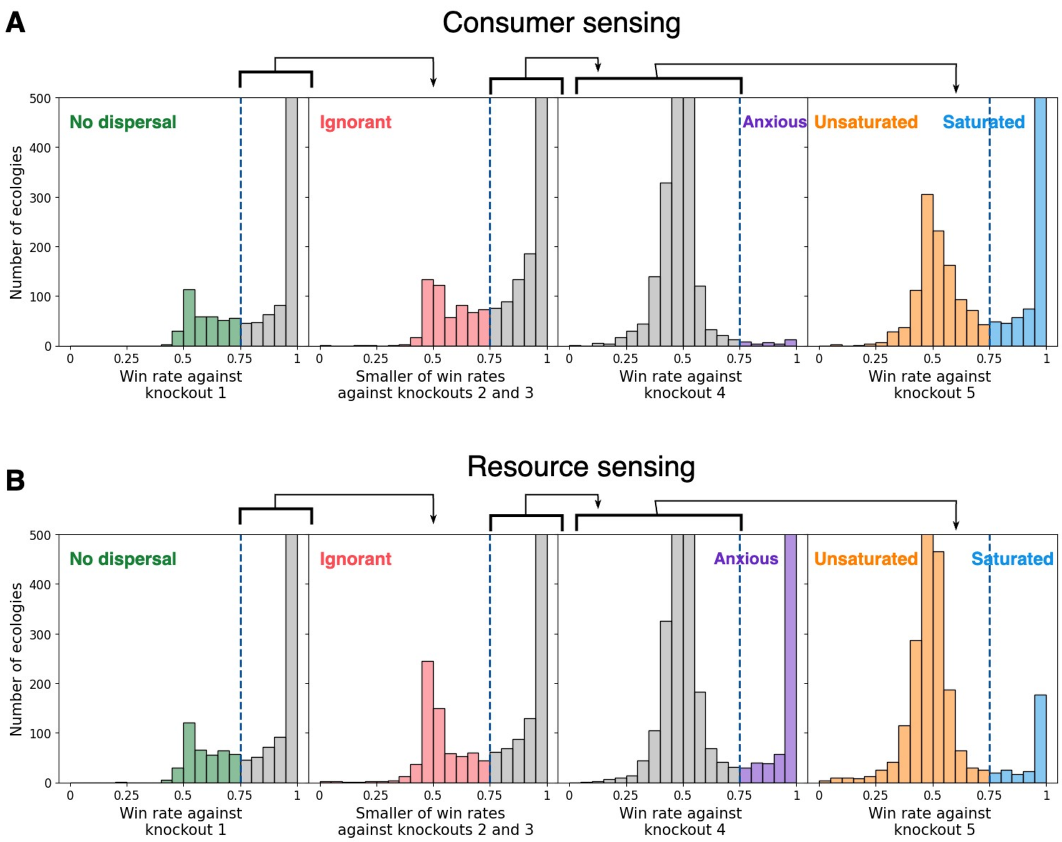

Figure S8: **Classification of evolved strategies into five classes.** Each plot shows the histogram of win rates of evolved strategies against knockouts. The vertical dashed lines are the thresholds determining the classes. Strategies in the colored section of a histogram are assigned to the indicated class. Strategies in the gray section of histogram are used in the next dichotomy (order is left to right, as indicated above the plots). Panel **A** shows the consumer sensing model, panel **B** shows the resource sensing model.

According to this definition, all five classes may appear in both the conspecifics sensing and the resource sensing models. However, the conspecifics sensing model presents only a handful strategies, which can be classified as anxious. Similarly, the resource sensing model features a few saturated strategies. The co-distribution of the strategies across classes in different models is presented in Table 3.

Table 3: The co-distribution of ecologies across classes in the conspecifics sensing and the resource sensing models. The table values show the number of ecologies that belong to the corresponding combination of classes in each model. The numbers above 250 are highlighted in bold. Asterisks indicate combinations, from which representative ecologies are sampled.

| | | Resource sensing model | | | | | |
|---|---|---|---|---|---|---|---|
| | | Non-viable | No dispersal | Ignorant | Anxious | Saturated | Unsaturated |
| Conspecifics sensing model | Non-viable | **9610** | 1 | 2 | **315** | 151 | **439** |
| | No dispersal | 4 | **351** | 10 | 0 | 0 | 2 |
| | Ignorant | 13 | 19 | **290*** | 13 | 11 | 210 |
| | Anxious | 1 | 0 | 2 | 4 | 5 | 24 |
| | Saturated | 29 | 18 | 135 | **359*** | 62 | **426*** |
| | Unsaturated | 32 | 10 | 232 | 34 | 31 | **756*** |

We select four representative ecologies to illustrate the most common combination of dispersal decision classes across the two informed dispersal models, assuming an ecology to be viable in all three models (including uninformed) but skipping the case of ecologies promoting no-dispersal in both models, see the ecological parameters in Table 4. Ecology 1 promotes ignorant strategies in both concpecifics and resource sensing models. Ecology 2 promotes a saturating strategy in the conspecifics sensing model and an anxious strategy in the resource sensing model. Ecology 3 promotes a saturating strategy in the conspecifics sensing model and an unsaturated strategy in the resource sensing model. Ecology 4 promotes unsaturated strategies in both concpecifics and resource sensing models.

Table 4: Ecological parameters of the four representative strategies.

| | $\omega_r$ | $K_r$ | $\omega_c$ | $h_c$ | $d_c$ | $E_r$ | Conspecifics sens. | Resource sens. |
|---|---|---|---|---|---|---|---|---|
| Ecology 1 | 1.261 | 1182 | 0.002463 | 0.3573 | 1.0 | 0.1 | Ignorant | Ignorant |
| Ecology 2 | 3.089 | 877 | 0.01771 | 0.4309 | 1.0 | 0.1 | Saturated | Anxious |
| Ecology 3 | 0.2536 | 1869 | 0.002546 | 0.02213 | 1.0 | 0.1 | Saturated | Unsaturated |
| Ecology 4 | 0.1446 | 2027 | 0.001734 | 0.2219 | 1.0 | 0.1 | Unsaturated | Unsaturated |

### 7 Resource-consumer dynamics in representative ecologies

In this section, we show the population dynamics in a course of ecological dynamics within one patch of the meta-population, see Fig. S9. These figures present complementary views on the within-patch dynamics. Note the presence of the giving-up density, emerging only in the resource sensing unsaturated strategy (the one matching closely to the MVT set up and predictions). No other representative ecologies demonstrate such a behavior.

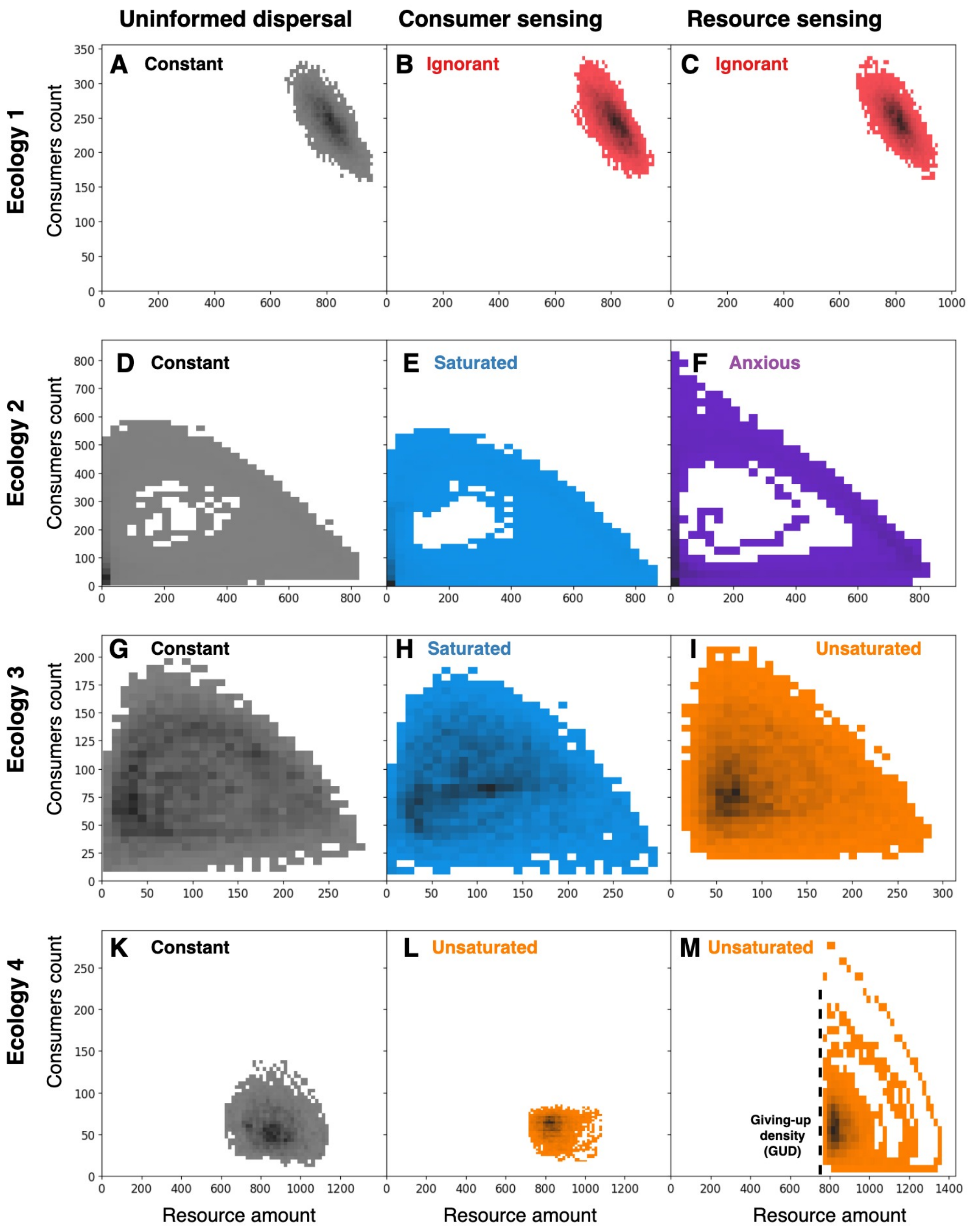

Figure S9: **Resource-consumer dynamics across models and ecologies.** Each panel shows a density of consumer/resource states in the course of a long (10000 simulation steps) dynamics in a single patch of a meta-community: darker shades indicate more likely states, white color indicates states never observed. The ecologies and strategies are the same as in the main text. Panel **M** (resource sensing unsaturated) features an emergence of a giving-up density – maximal resource population, at which a consumer states in the population.

### 8 Comparison of within-patch dynamics between informed and uninformed sensing models

We can infer the impact of the informed dispersal by comparing the population dynamics occurring in the same ecology in an informed and the uninformed sensing models. For this analysis, we only take into account ecologies, which present viable populations in both informed and uninformed models (the list of such ecologies depends on whether the informed model is the conspecifics sensing or the resource sensing). First, we characterize the difference in dynamics using the ratio of median number of consumers and resource observed in a patch in the course of simulation:

$$R = \frac{\text{median}(N_{\text{informed}})}{\text{median}(N_{\text{uninformed}})}, \tag{54}$$

where $N$ could be either the number of consumers in a patch $N_c$ or amount of resource in a patch $N_r$, and the informed model could be either conspecifics or resource sensing.

The distribution of medians ratios are very close to one for both informed models and for both species, see Fig. S10.

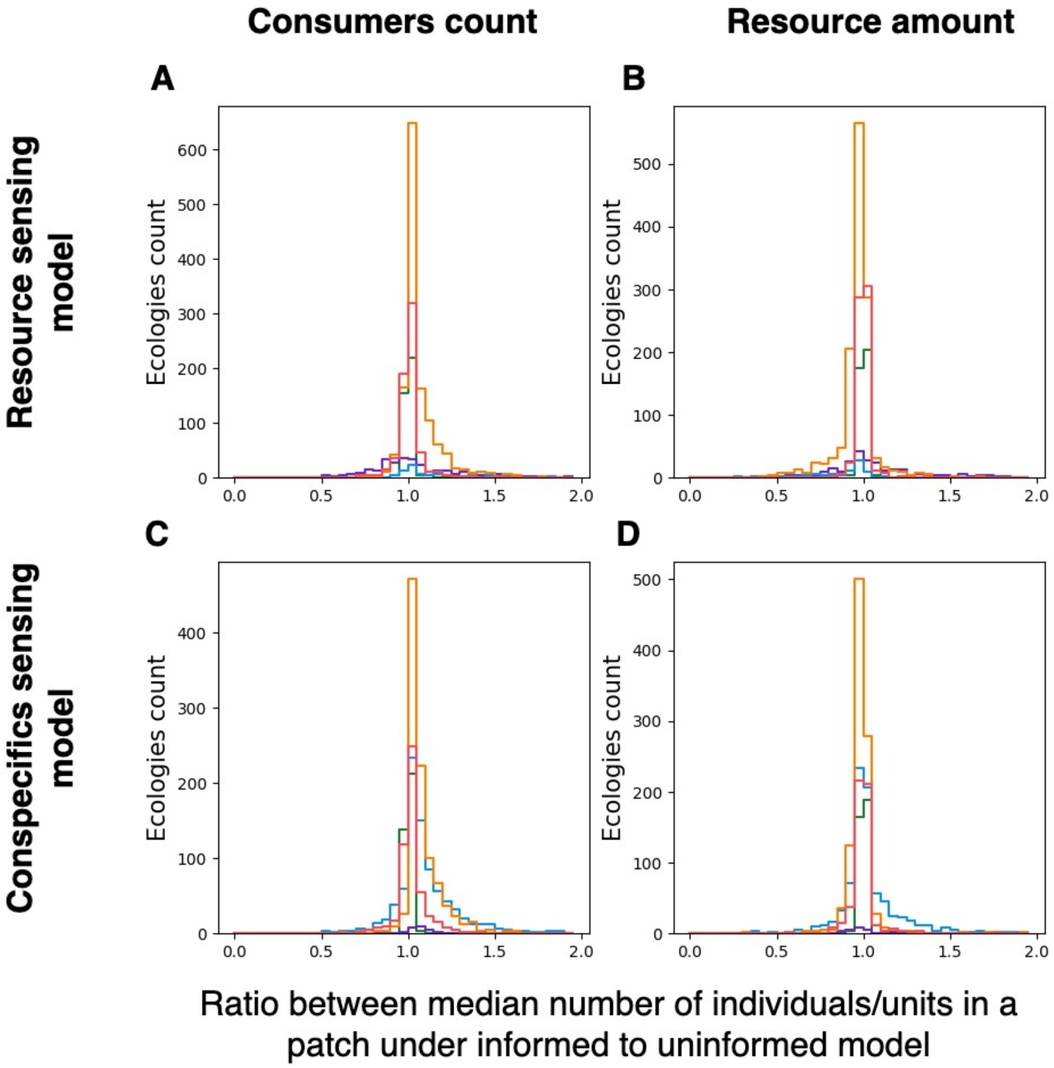

Figure S10: **Informed dispersal has little impact on the long-term median patch state.** Panels show the distributions of median ratios of the consumer count (panels **A, C**) and the resource amount (Panels **B,D**) between informed and uninformed model separated by class (color). Panel **A** - ratio of medians of the consumer counts in the resource sensing model. Panel **B** - ratio of medians of the resource amount in the resource sensing model. Panel **C** - ratio of medians of the consumer count in the conspecifics sensing model. Panel **D** - ratio of medians of the resource amount in the conspecifics sensing model. Only the ecologies viable in both uninformed and the referenced informed sensing models are considered.

Next, we characterize the difference in dynamics using the ratio of ranges of consumers and resource numbers observed in a patch in the course of simulation:

$$R = \frac{\max(N_{\text{informed}}) - \min(N_{\text{informed}})}{\max(N_{\text{uninformed}}) - \min(N_{\text{uninformed}})}, \tag{55}$$

where $N$ could be either the number of consumers in a patch $N_c$ or amount of resource in a patch $N_r$, and the informed model could be either conspecifics or resource sensing. The ratio value below one indicates a dampening of exhibited variation in population size, while the ratio above one indicates an amplification of said variation. The distribution of these ratios for different populations and sensing models are shown at Fig. S11.

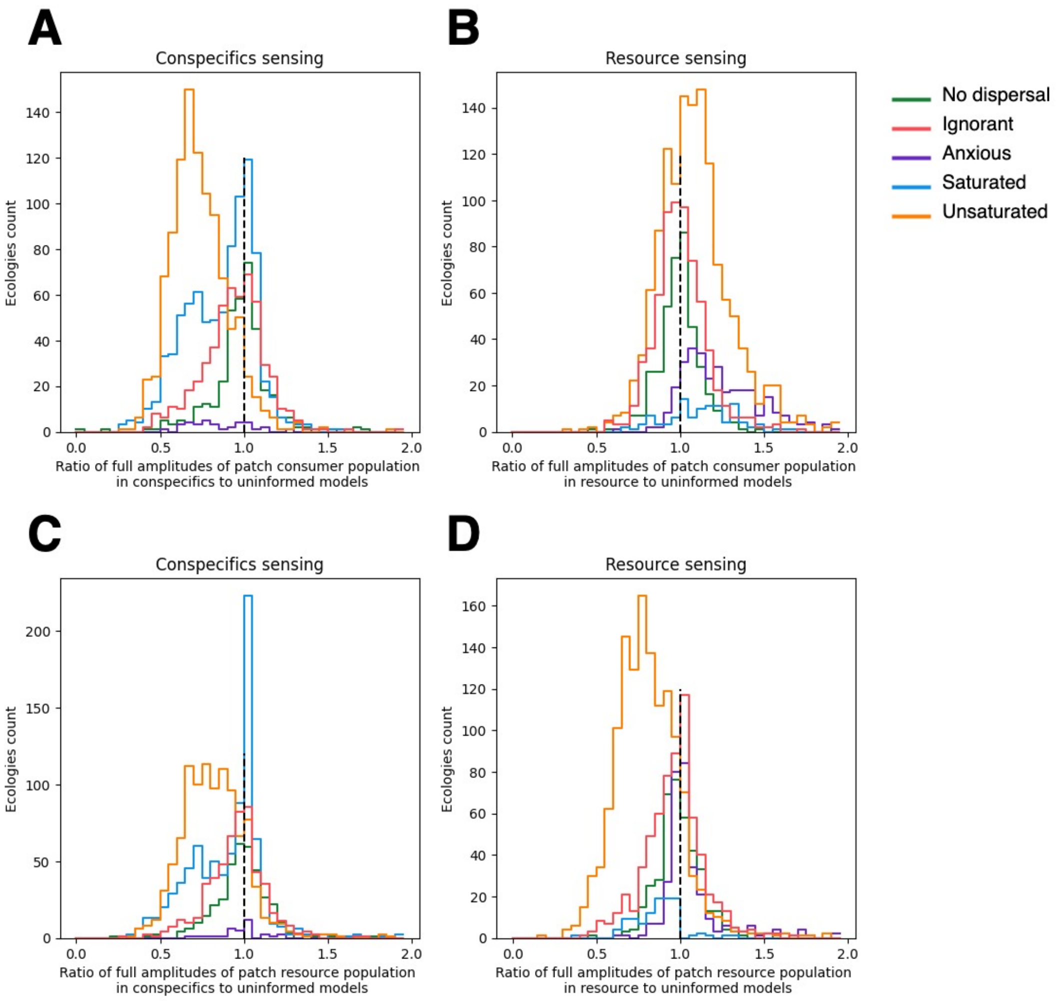

Figure S11: **Different classes have different impact on the within-patch dynamics** Panels show the distributions of range ratios of the consumer count (panels **A, C**) and the resource amount (Panels **B,D**) between informed and uninformed model separated by class (color). Panel **A** - ratio of ranges of consumer counts in the conspecifics sensing model. Panel **B** - ratio of ranges of consumer counts in the resource sensing model. Panel **C** - ratio of ranges of the resource amount in the conspecifics sensing model. Panel **D** - ratio of ranges of the resource amount in the resource sensing model. Only the ecologies viable in both uninformed and the referenced informed sensing models are considered.

The model is stochastic, and hence all metrics are inherently noisy. However, the *No-dispersal* and *Ignorant* classes show us the scale of this noise. These strategies are nearly identical between informed and non-informed models. Therefore, deviation of range ratios from one observed in these classes can be attributed to stochastic differences between different replicates of (nearly) identical dynamics. Hence, below, we only consider notable deviations from distributions exhibited by these two classes.

In the conspecifics sensing model, the *Unsaturated* class dampens oscillations in both consumer

(Fig. S11A) and resource dynamics (Fig. S11C) and the example of such dynamics is given by the representaive ecology 4 at Fig. S9L. Here, the rapid feedback loop between consumer population size and dispersal rate suppresses the fluctuations, as explained in details in the main text.

The *Saturated* strategies in this model exhibit bi-modal distributions of ranges ratio for both consumers (Fig. S11A) and resource (Fig. S11C): in some ecologies the range of population variation is dampened in conspecific sensing compared to uninformed one (similar to the *Unsaturated* class), while in others the range of variation is not affected by the informed model. Mapping these ecologies onto the ecological space, reveals that the ecologies dampening variation (blue shades on Fig. S12) have small consumer population, while ecologies preserving them (black shades on Fig. S12) tend to have a larger consumer population. While the *Saturated* strategies are associated with limit cycles, in ecologies with small population size, the main source of population size variation are the fluctuations around the limit cycle. These fluctuations can be efficiently suppressed by the conspecifics sensing strategy, as was shown for *Unsaturated* class. However, with an increase in population size, the impact of fluctuations becomes relatively smaller, compared to the amplitude of the limit cycle itself. This inherent source of range variation cannot be suppressed by dispersal, and when dominant, preserves the ranges ratio between conspecifics sensing and uninformed models.

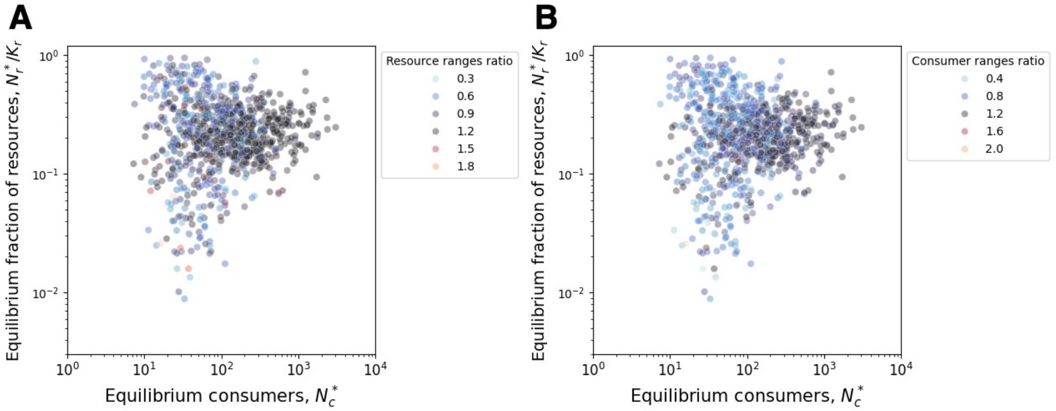

Figure S12: **Variation range is suppressed by *Saturated* strategies in small-size populations and preserved in large-size populations** Panels represent the separation of two peaks of the bi-modal ranges ratio distribution of the *Saturated* strategies under consumer-sensing model in ecological space. Each point represents an ecology promoting a *Saturated* strategy under the conspecifics sensing model and viable under the uninformed model. The color of the point shows the ratio of ranges of resource in a patch (panel **A**) in two models or the ratio of ranges of the consumers in a patch (panel **B**). Blue shades indicates a suppression under conspecifics sensing, black shades - preservation, red shades - amplification.

We also address the high number of ecologies, which feature the resource ranges ratio of nearly 1 in this class. These are the ecologies, in which the resource population occasionally fall to very small numbers but can also recover up to the carrying capacity ($N_r = K_r$) in both uninformed and conspecifics sensing models. Thus, the range of resource amount in these ecologies is close to $K_r$ in both models, which turns the ranges ratio to be nearly one with very little noise possible.

In the resource sensing model, the *Unsaturated* class tends to dampen the variation of the resource amount (Fig. S11D) but tends to amplify the variation of the consumers numbers (Fig. S11B). An elevated dispersal from the resource depleting patch does not allow the resource population to fall to low numbers, as demonstrated by representative ecology 4 on Fig. S9M. Hence, the range of the resource population size is smaller under the informed dispersal. At the same time, the same mechanism is responsible for the amplification of the range of consumer population size. Elevated dispersal persists until the resource population is recovered but by the moment it happens, the size of consumer population falls lower than it would be in the uninformed model. In addition, under the resource-rich conditions, a negligible outflow of migrants does not balance their inflow, rising the peak population size above available for uninformed model. Altogether, the higher peak and the lower bottom of consumer counts in the resource sensing model amplify its variation comapred to the uninformed model. As a result, *Unsaturated* strategies in the resource sensing model have an opposite effect on the dynamics of the resource (dampening the variation) and consumers (amplification the variation).

*Anxious* strategies have a little impact on the resource size range (see Fig. S11D). Since the *Anxious* strategies emerge under limit cycles with large population size, the variation in the resource amount is driven by the amplitude of the limit cycle, on which the dispersal has a negligible impact. At the same time, *Anxious* strategies amplify the variation of the number of consumers (see Fig. S11B). The amplification of the consumers number range can be attributed to the higher peak population size, achieved just before the population collapse (see Fig. S9F). At around the peak of consumer population, the lagging patch experiences an imbalance of migration flows. The outgoing migrants depart at the base-line moderate rate, since the local resource is not depleted yet. However, the incoming migrants come from patches ahead in the limit cycle, where the resource is depleted and the dispersal rate from there is very high. This net positive migration flow pushes the peak population of the lagging patch above the values available to uninformed model (where dispersal rates are the same in all patches), conspecifics sensing model (where the imbalance of the migration flows occurs at a moderate overpopulation level).
